# golgi: an open-source graphical platform for image-to-recruitment modeling of peripheral nerve stimulation

**DOI:** 10.64898/2026.07.10.737529

**Authors:** David Lung, Yuting Jia, Roland Blumer, Lukas Reissig, Lydia M. Zopf, Patrick Heimel, Christoph Kraus, Andrea Moro, Matteo Fachino, Max Haberbusch

## Abstract

Computational models of peripheral nerve stimulation—coupling finite-element bioelectric fields to biophysical axon models—have become essential for designing electrodes and waveforms for neuromodulation therapies. Yet the established open tools are code-first and assume substantial modeling expertise, and several depend on commercial finite-element solvers, placing realistic nerve modeling out of reach for many experimentalists and clinicians. We present *golgi*, an open-source platform that takes a peripheral nerve from image to stimulated fiber population through a single graphical interface, with an equivalent scriptable Python API and command-line interface for batch studies. *golgi* integrates the full pipeline: image segmentation (or import of surfaces or masks), automated multi-region tetrahedral meshing, anisotropic finite-element solution of the extracellular field with explicit perineurium contact impedance, generation of realistic fiber populations and their three-dimensional trajectories (straight, or curved via a quasi-static streamline solver), and biophysical activation thresholds through interchangeable NEURON and a GPU-accelerated surrogate backend. We demonstrate *golgi* on extruded multifascicular swine and human cervical vagus nerves and on real three-dimensional, branching human and rabbit vagus nerves reconstructed from micro-computed tomography. It reproduces the physiological fiber-diameter recruitment order, quantifies fascicular selectivity and current steering with a multi-contact cuff electrode, and resolves anatomically defined nerve branches. Using this branch resolution, we find that selectively engaging a vagal cardiac branch from a proximal cuff depends on anatomy. In the rabbit, whose cardiac fibers are predominantly small and whose superior cardiac branch forms a discrete, spatially segregated tract, current steering isolates even its small cardiac (B-type) fibers; in the human cervical vagus only the large myelinated fibers are separable, because the high thresholds of the small cardiac fibers force stimulus currents that also recruit off-target fibers. Every study can be exported as an integrity-hashed, self-contained bundle whose finite-element-to-recruitment provenance is verifiable byte-for-byte with a single command—a reproducibility guarantee absent from existing tools. By combining non-specialist usability, anatomical realism, and verifiable reproducibility in one open package, *golgi* lowers the barrier to in-silico peripheral nerve stimulation modeling. *golgi* is freely available as open-source software.

**Author summary:** Electrical stimulation of peripheral nerves treats a growing range of conditions, from epilepsy to inflammatory and cardiovascular disease. Deciding where to place an electrode and how to shape the stimulus increasingly relies on computer models that combine the electric field around the electrode with detailed models of how individual nerve fibers respond. We found that existing software for this is powerful but primarily designed for expert modelers: it generally requires programming, substantial modeling expertise, and sometimes expensive commercial software, which can limit its adoption by experimentalists and clinicians. We built *golgi* to remove that barrier. With *golgi*, a user can go from a nerve image all the way to predicted fiber recruitment through a single point-and-click interface, while advanced users keep full scripting control. *golgi* builds anatomically realistic nerve models, simulates how different fiber types and fascicles are recruited, and lets users compare electrode designs. Using *golgi*, we also found that whether a small but clinically important nerve branch—such as the cardiac branch of the vagus nerve—can be selectively stimulated depends on its anatomy. In a rabbit nerve, where this branch forms a discrete, spatially separated bundle and its fibers are mostly small, even its small fibers can be targeted from a cuff on the main trunk; in the human, only the large fibers can be reached selectively, because the small cardiac fibers are harder to excite and the stronger currents needed to reach them also activate off-target fibers. Critically, every result can be packaged so that anyone else can verify it reproduces exactly—making peripheral nerve stimulation models easier to build, share, and trust.

## Introduction

Electrical stimulation of peripheral nerves underlies a rapidly expanding set of neuromodulation therapies, including vagus nerve stimulation for epilepsy [1], depression [2], and inflammatory [3] and cardiovascular [4] disease. The therapeutic effect, and the off-target side effects that limit it, are determined by which nerve fibers a given electrode and waveform recruit—a function of the fiber’s diameter and type, its position within the fascicular anatomy, and the spatial distribution of the extracellular potential the electrode produces [5–7]. Because these dependencies are difficult to probe experimentally, computational models have become a central tool for rationally designing electrodes, contact configurations, and stimulus parameters [8, 9].

The standard modeling approach couples a volume-conductor description of the tissue—most often solved by the finite-element method (FEM)—to biophysical cable models of individual axons [10, 11]. The extracellular potential computed along each fiber drives a multi-compartment membrane model, and the second spatial derivative of that potential, the activating function, predicts where the fiber is depolarized [12]. Validated myelinated [11, 13] and unmyelinated [14–19] fiber models, combined with realistic fascicular geometry, allow recruitment and selectivity to be estimated in silico.

Several open frameworks now implement variants of this pipeline. ASCENT couples automated nerve-geometry construction to a commercial FEM solver and NEURON fiber models [8]; NRV provides a Python framework for in-silico evaluation of stimulation strategies [9]; PyPNS [20] and ViNERS [21] target related niches. These tools have substantially advanced reproducible peripheral nerve modeling. However, they share two practical barriers. First, they are code-first: building and running a study requires programming and detailed modeling expertise, and ASCENT additionally requires a commercial FEM solver, which together exclude many of the experimental and clinical users who would benefit most. Second, despite being scriptable, none packages a complete study—geometry, mesh, field solution, fiber population, and simulation results—into a single artifact whose end-to-end provenance can be independently verified, so reproducing a published result in practice remains effortful and error-prone.

Here we present *golgi* —an homage to Camillo Golgi, whose silver stain first revealed individual nerve cells in their entirety—an open-source platform designed to occupy the gap between accessibility and realism. *golgi* exposes the entire image-to-recruitment pipeline through a graphical interface usable without programming, while mirroring every operation in a scriptable Python API and command-line interface for batch and high-performance use. It builds anatomically realistic, multifascicular nerve models—directly from segmented images, imported surfaces, or label masks—meshes and solves them with a fully open FEM stack, generates realistic fiber populations with straight or physically curved three-dimensional trajectories, and computes biophysical thresholds through interchangeable simulation backends. Uniquely, *golgi* exports each study as an integrity-hashed, self-contained bundle whose provenance is verifiable byte-for-byte. We describe *golgi*’s design and demonstrate it on extruded multifascicular swine and human cervical vagus nerves and real three-dimensional branching human and rabbit vagus nerves reconstructed from micro-computed tomography images—using the branching nerves to ask how selectively a vagal cardiac branch can be engaged from a proximal cuff.

## Design and implementation

### Overview and access modes

*golgi* is organized as a unified computational pipeline that begins with image or geometric data, proceeds through multi-region mesh generation and finite-element field construction, generates fiber populations and trajectories for fiber simulation, and concludes with downstream analysis (Fig 1). The same pipeline is exposed through three complementary interfaces: an interactive browser-based graphical user interface (GUI) for users who prefer point-and-click operation; a headless Python API (golgi.Study) that mirrors every GUI action for scripting and batch studies; and a command-line interface for cluster and continuous-integration use. This design allows a non-programming user to complete a full study graphically (S1 Fig) while enabling an advanced user to script identical studies or offload them to a high-performance computing scheduler, with results that are interchangeable across modes.

**Fig 1.**
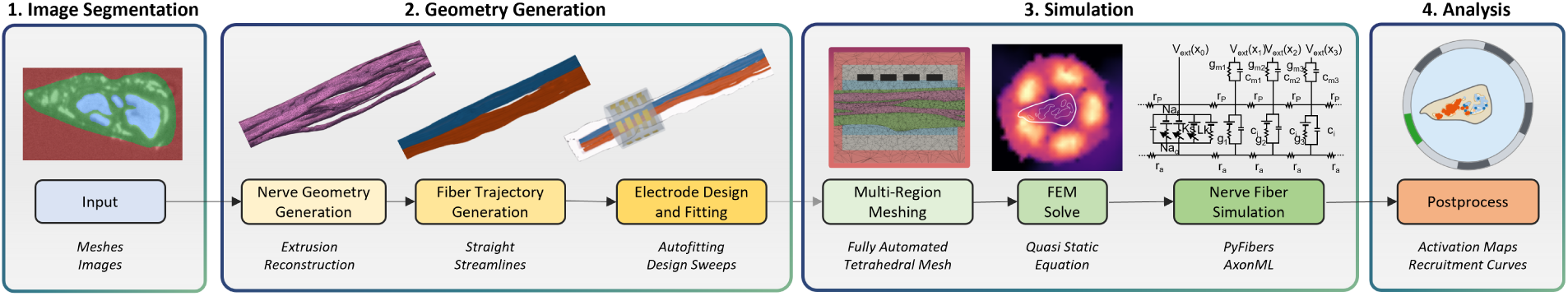
*golgi* spans the full peripheral nerve stimulation modeling pipeline through a no-code interface. End-to-end pipeline. From an input image, surface, or mask, *golgi* reconstructs the nerve geometry, generates fiber trajectories (straight extrusion or quasi-static streamlines), designs and fits electrodes, builds multi-region tetrahedral meshes, solves the quasi-static finite-element field, simulates nerve fibers through a choice of backends (NEURON via PyFibers [22], or the GPU-accelerated surrogate AxonML [23]), and post-processes activation maps and recruitment curves.

While *golgi* is designed around its graphical interface for accessibility, the same image-to-recruitment pipeline (Fig 1) is fully scriptable and headless: the golgi.Study Python API mirrors every graphical step and a command-line interface exposes the same operations, so an identical study can be reproduced, batched, or offloaded to a high-performance scheduler without ever opening the GUI.

**Listing 1.**
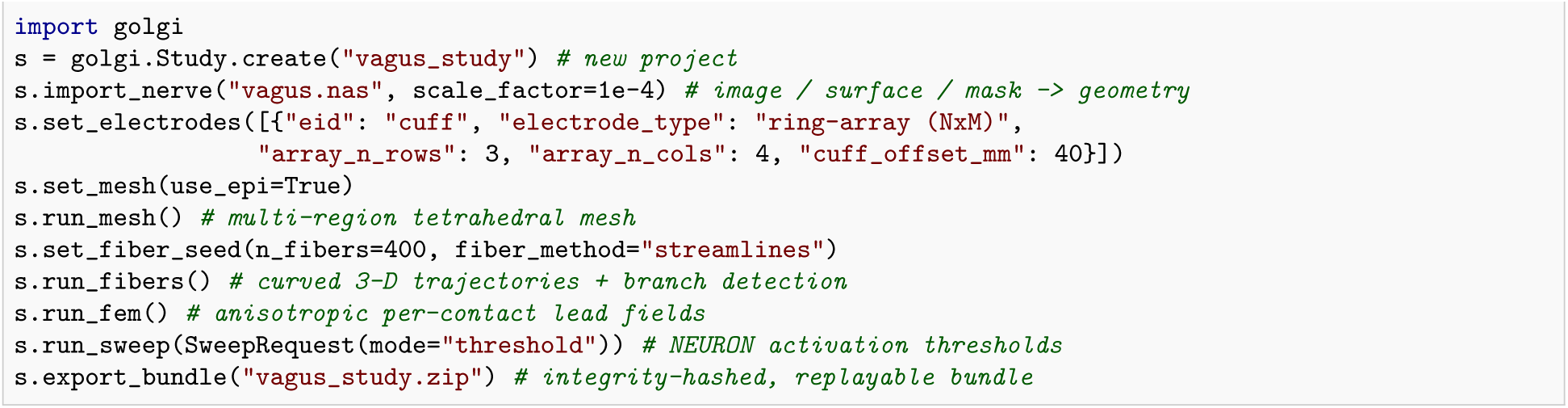
The Fig 1 pipeline expressed through the headless golgi.Study API.

### Geometry, segmentation, and meshing

Nerve geometry can be created three ways: by segmenting a volumetric image (e.g., from a micro-computed tomography stack) with an integrated promptable segmentation engine guided by point and box prompts, followed by interactive mask refinement (per-class reassignment, brush add/erase) [24, 25]; by importing a surface mesh (STL/NAS/OBJ); or by importing label masks (e.g., from segmented histology slices). Segmented or imported nerve cross-sections are reconstructed into a three-dimensional nerve (Fig 2A), either by extrusion of a representative cross-section or, for genuine three-dimensional samples, from the full stack. *golgi* then builds a multi-region tetrahedral mesh (TetGen/Gmsh [26, 27]) comprising endoneurium, epineurium, the electrode and its insulating cuff, and the surrounding bath, with mesh-quality visualization (Fig 3A). The perineurium—the high-resistance fascicle sheath—is not meshed as a separate volume but instead enters the model as a thin-layer contact-impedance interface at the endoneurium–epineurium boundary.

**Fig 2.**
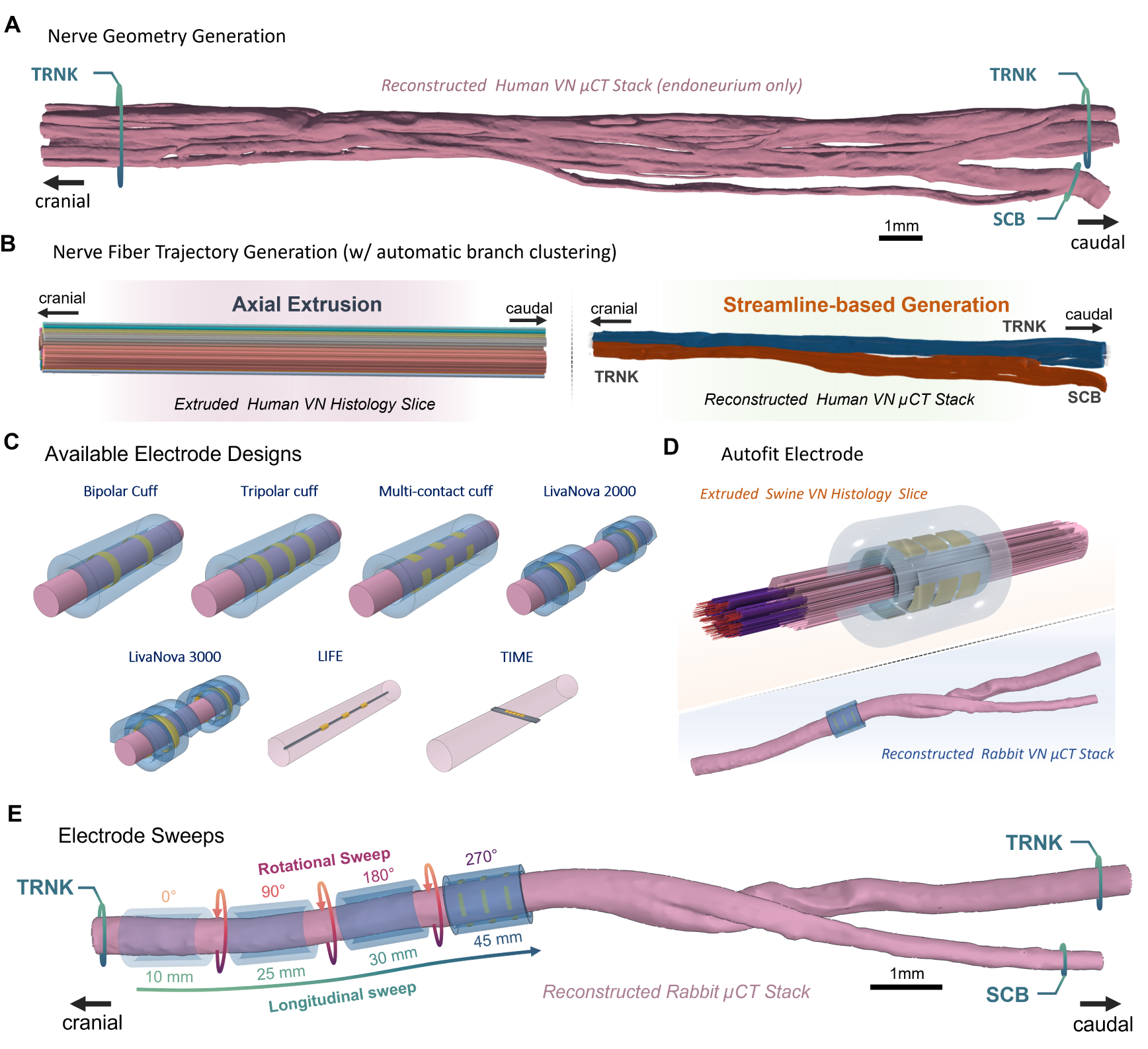
Geometry reconstruction, fiber trajectories, and electrode design. **A:** Nerve-geometry generation—a real three-dimensional human cervical vagus reconstructed from a micro-computed tomography stack (endoneurium shown), with the common trunk (TRNK) and superior cardiac branch (SCB). **B:** Fiber-trajectory generation with automatic branch clustering—straight axial extrusion (left) versus quasi-static streamline integration that follows the true endoneurial course through the bifurcation (right). **C:** Library of parameterized electrode designs—bipolar, tripolar, and multi-contact cuffs, LivaNova-style helical cuffs, and intrafascicular electrodes—longitudinal intrafascicular electrodes (LIFEs) and transverse intrafascicular multichannel electrodes (TIMEs). **D:** Automatic fitting of an electrode to the local nerve axis, shown on an extruded swine and a reconstructed rabbit nerve. **E:** Rotational and longitudinal cuff-position sweeps, shown on a reconstructed rabbit nerve.

**Fig 3.**
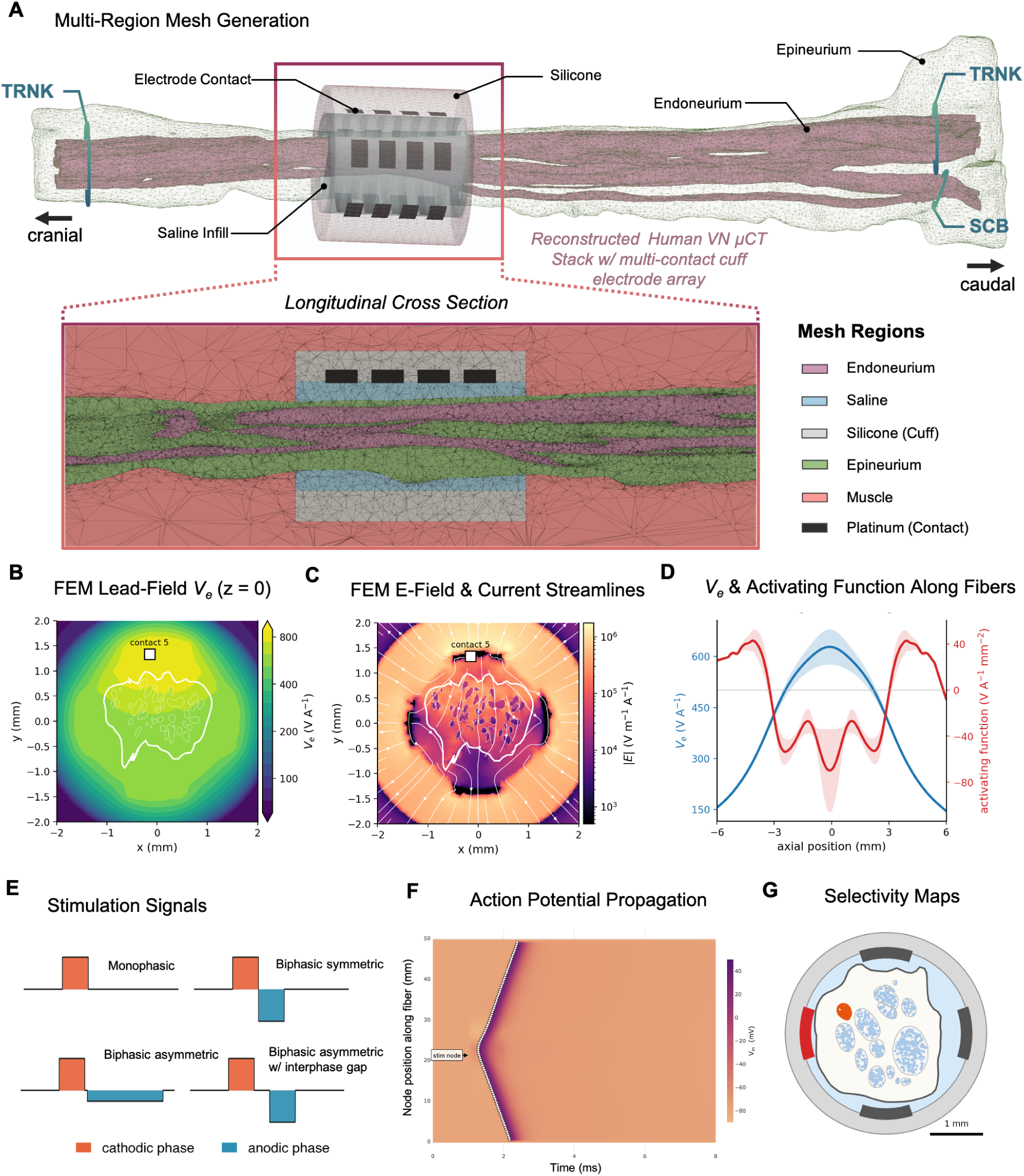
Multi-region meshing, finite-element field, and fiber drive. **A:** Multi-region tetrahedral mesh (longitudinal cross-section) of a reconstructed human cervical vagus in a multi-contact cuff, coloured by tissue region: endoneurium, saline, silicone cuff, epineurium, muscle, and platinum contacts. **B:** Finite-element per-contact lead-field extracellular potential *V_e_* on a cuff cross-section (central contact, *z* = 0), with nerve and fascicle boundaries overlaid. **C:** Corresponding electric-field magnitude with in-plane current streamlines. **D:** Mean ± s.d. of the smoothed *V_e_* (blue) and its activating function d^2^*V_e_/*d*s*^2^ (red) sampled along the fiber population, linking the volumetric field (b,c) to the per-fiber drive. **E:** Configurable stimulation waveforms (monophasic; biphasic symmetric; biphasic asymmetric, with optional interphase gap). **F:** Evoked action-potential propagation along a fiber (membrane potential in space and time). **G:** Fascicular selectivity map—per-fascicle activation across the cross-section for a chosen contact.

### Electrode design and finite-element field

An interactive electrode designer lets the user place and parameterize electrodes—from bipolar and multi-contact cuffs to intrafascicular electrodes (Fig 2C)—and automatically fits the electrode to the local longitudinal axis of the nerve (Fig 2D), with the possibility to also perform rotational and longitudinal cuff-position sweeps (Fig 2E). *golgi* solves the quasi-static [28] volume-conductor problem with the open-source FEniCSx/DOLFINx finite-element library [29], discretizing the field with continuous piecewise-linear (P1) Lagrange tetrahedral elements. On the domain Ω (endoneurium, epineurium, electrode contacts, insulating cuff, and surrounding bath/muscle) it solves

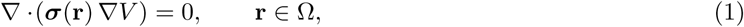

where ***σ*** is the (anisotropic, *z*-longitudinal in the endoneurium and muscle) conductivity tensor (Table 1). Because a cuff electrode does not appose the nerve perfectly, the gap between the cuff and the epineurium is modeled as a conductive saline bath, which provides the electrical path from the contacts to the nerve surface, as in ASCENT [8]. The bath outer boundary is grounded (*V* = 0); each electrode contact injects a prescribed current and the remaining cuff surface is insulating:

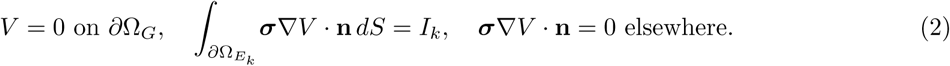

**Table 1.**
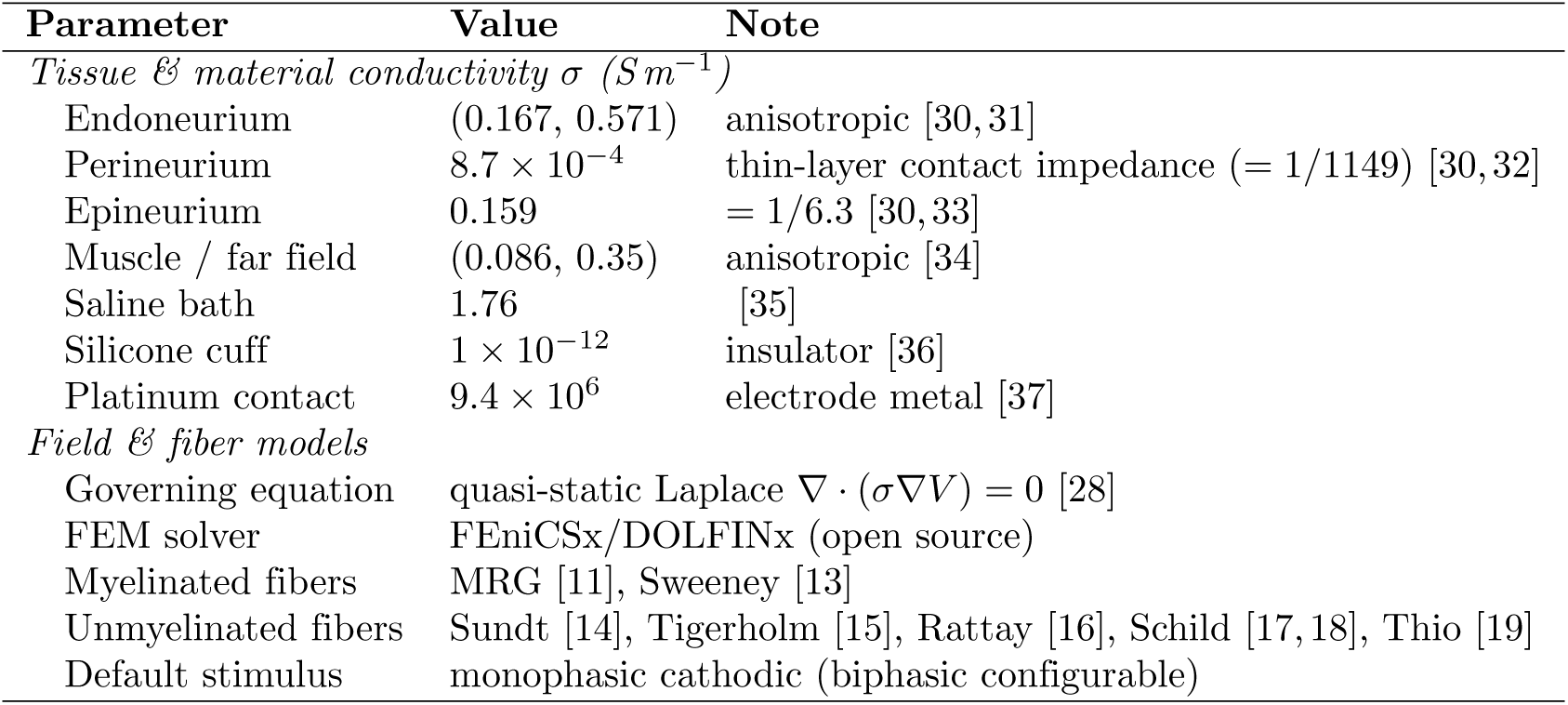
Tissue, material, and model parameters used by *golgi*. Conductivities follow the finite-element parameter set of Pelot et al. [30]; anisotropic tissues are listed as (transverse, longitudinal) along the nerve/muscle-fiber axis.

The perineurium is not meshed as a volume but enters as a thin-layer contact-impedance interface Γ between the endoneurial and extra-fascicular domains, with a sheet resistance *R_s_* = *t_p_/σ_p_* that induces a potential jump for the normal current density,

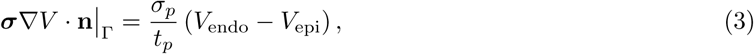

implemented as a two-field Robin coupling of the endoneurial and extra-fascicular subproblems [30]. Because the medium is linear, the field obeys superposition: a per-contact lead field *ϕ_k_* is obtained once from a unit-current solve and reused, so the potential of any multi-contact montage {*I_k_*} is assembled without re-solving,

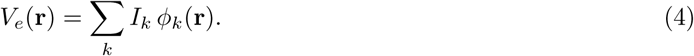

The block system is factorized once with a sparse direct solver (MUMPS) and back-substituted for each contact, so arbitrary multi-contact configurations and current-steering combinations are inexpensive to evaluate; the open FEniCSx solver is verified against the analytic point-source potential, cross-validated against a commercial finite-element reference (COMSOL), and checked against classical fiber-model scaling laws (Fig 4A,B, S8 Fig, S15 Fig). The resulting extracellular potential and electric field (Fig 3B,C) are sampled along each fiber to yield the potential profile and its activating function (Fig 3D), the drive for the downstream fiber models, which are stimulated by configurable stimulation waveforms (Fig 3E); a fiber driven above threshold fires a propagating action potential (Fig 3F), and aggregating activation across the population yields a per-fascicle selectivity map for a chosen contact (Fig 3G).

**Fig 4.**
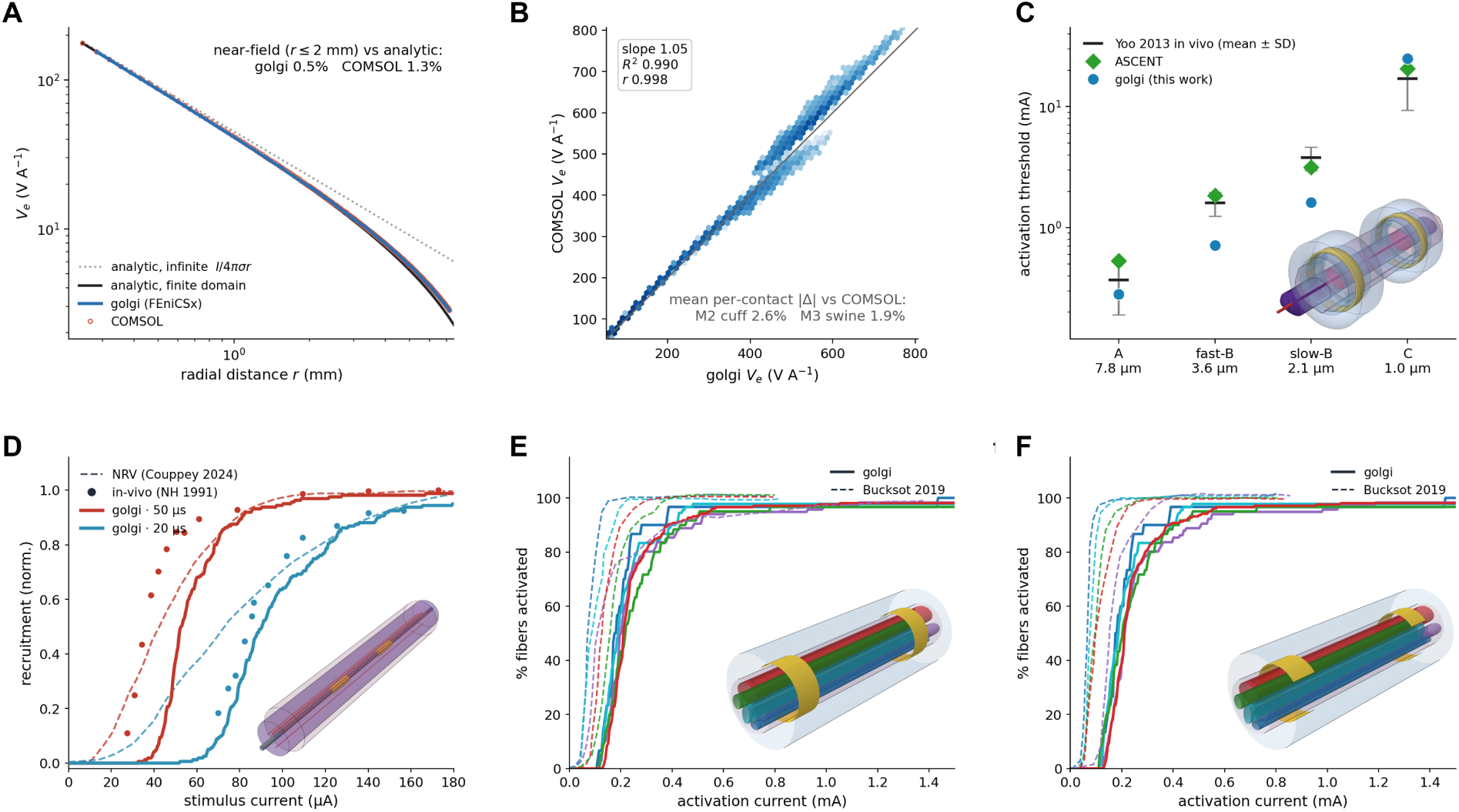
Validation of *golgi* against a commercial finite-element solver and against independent experimental and modeling benchmarks. Panels **A** and **B** test the open FEniCSx field solver; panels **C**–**F** test the complete image-to-recruitment pipeline. **A:** Extracellular potential *V_e_*(*r*) around a current monopole in a grounded saline cylinder, computed by *golgi* (FEniCSx) and by COMSOL and compared with the analytic solution *<u>^I^</u>* ^1^ − <u>^1^</u> (the infinite-medium term *I/*4*πσr* is shown dotted). Both solvers reproduce the analytic potential to a near-field (*r* ≤ 2 mm) mean error of 0.5 % (*golgi*) and 1.3 % (COMSOL). **B:** Per-contact lead fields computed by *golgi* and by COMSOL for a reconstructed swine cervical vagus nerve (40 fascicles), compared point for point across 12 contacts and ∼10^5^ sample positions (density scatter; the line marks identity). The mean per-contact difference is 1.9 % for the swine nerve and 2.6 % for an idealized cuff model (*R*^2^ ≥ 0.98). Full per-model comparisons are given in S15 Fig. C: *In vivo* dog cervical VNS activation thresholds for A, fast-B, slow-B, and C fibers (300 µs monophasic cathodic pulse): experimental data (Yoo et al. 2013; mean ± SD), ASCENT’s COMSOL-based predictions, and *golgi*. The model was built by extruding ASCENT’s nerve and fascicle masks through the *golgi* pipeline—a 1.7 mm fascicle within a 3.27 mm nerve, with a perineurium contact-impedance sheet, a separated LivaNova-style bipolar cuff, and surrounding muscle (inset: cel-shaded rendering, with epineurium in rose, the endoneurial fascicle in purple, the centroid fiber in red, and gold ring contacts). *golgi* reproduces the correct recruitment order; its A-fiber threshold lies within the in-vivo SD band and its C-fiber at the upper edge of the in-vivo range, comparable to ASCENT, whereas the small B-fibers (below the MRG calibration range) are underestimated roughly two-fold. **D:** Recruitment versus stimulus current for a longitudinal intrafascicular electrode (LIFE) in a monofascicular nerve at 20 and 50 µs (inset): *golgi* (solid), NRV (dashed; Couppey et al. 2024), and *in vivo* data (points; Nannini & Horch 1991). *golgi* falls within the in-vivo and NRV bands, with a strength–duration rate ratio of 2.0 (in vivo 2.1; NRV 2.4). **E,F:** Per-fascicle recruitment (percentage of fibers activated versus current) in a five-fascicle nerve under a circumferential bipolar cuff: *golgi* (solid) and the digitized model of Bucksot et al. 2019 (dashed), for the original contact (**e**) and the same 270*^◦^* contact rotated by 180*^◦^* (**f**). Fascicle colours match the setup rendering (inset). *golgi* reproduces the published thresholds and the near orientation-independence of circumferential recruitment.

### Fiber populations and trajectories

*golgi* allows the user to populate each fascicle or monofascicular nerve with fibers drawn from curated species-and nerve-specific diameter and fiber-type statistics—sourced from published vagal morphometry where available (each in-application preset carrying its own citation) or set directly by the user—mixing myelinated (A- and B-type) and unmyelinated (C-type) fibers in the desired proportions. Fiber trajectories are generated either as straight axial extrusions or, for three-dimensional and branching geometries, as smooth curved paths obtained by integrating streamlines of a quasi-static (Laplace) field solved within the endoneurial domain, following the approach of Marshall et al. [38]—capturing the realistic course of fibers through curved and bifurcating nerves (Fig 2B). For a branching nerve, these streamlines diverge at the bifurcation and terminate in the separate distal cross-sections of the daughter branches; *golgi* clusters the fibers by the branch ending they reach—which requires only that the clustering threshold be smaller than the inter-branch separation—and thereby labels every fiber by its destination branch. This per-fiber branch assignment is the prerequisite for branch-selective analysis: for each fiber passing under a proximally placed cuff it identifies whether the fiber belongs to a target branch (such as a cardiac branch) or to the off-target trunk continuation, so on-versus off-target recruitment can be quantified—an assignment that straight-fiber pipelines, which have no branch structure, cannot provide.

### Fiber simulation backends

Activation thresholds and responses are computed with biophysically detailed axon models. *golgi* exposes interchangeable backends behind a common interface: PyFibers [22, 39], which runs a library of validated myelinated (MRG [11], Sweeney [13]) and unmyelinated (Sundt [14], Tigerholm [15], Rattay [16], Schild [17, 18], and Thio autonomic/cutaneous [19]) axon models in the NEURON simulator [40] as the gold standard; and AxonML [23, 41], a GPU-accelerated surrogate for high-throughput sweeps. Thresholds are found by bisection on the amplitude of a cathodic (cathode-leading) monophasic pulse by default; biphasic and anode-leading waveforms are equally configurable. To avoid the non-physical “end-excitation” artifact—spurious activation at the truncated fiber terminals under long pulses—*golgi* tapers the extracellular drive to zero at the fiber ends, so that thresholds reflect genuine, centrally initiated action potentials.

### Reproducibility, provenance, and sharing

Every golgi study can be exported as a single self-contained bundle. The bundle’s manifest records the processing pipeline as a directed acyclic graph in which each stage feeds the next—from meshing through the field solution, fiber generation, and simulation to the final recruitment sweep—with a SHA-256 hash of every input and output file and of each stage, together with the exact software version and a frozen dependency list. A single golgi replay command re-hashes a received bundle and verifies, byte-for-byte, that it is intact and matches its manifest; a flight-recorder audit log captures the sequence of actions that produced it. This makes a complete image-to-recruitment study independently verifiable and shareable.

### Software implementation and requirements

*golgi* is open-source software implemented in Python and runs on Linux and macOS. It builds on the FEniCSx/DOLFINx finite element framework, NEURON for biophysical fiber simulation, and PyFibers for fiber trajectory handling; an optional GPU-accelerated backend (AxonML) can be enabled where compatible hardware is available. Beyond the terms of its AGPL-3.0-or-later license, *golgi* imposes no restrictions on use; however, because it links the copyleft mesh generators TetGen (AGPL-3.0, via the tetgen module) and Gmsh (GPLv2-or-later), redistributing *golgi* as part of a closed-source or commercial product additionally requires separate commercial licenses for TetGen (from WIAS) and Gmsh. The optional AxonML GPU backend is licensed separately by Duke University for non-commercial academic use and is not bundled with *golgi*.

## Results

We illustrate *golgi*’s capabilities through a sequence of analyses on porcine, human, and rabbit nerve samples, all produced through the same pipeline that drives the graphical interface. Each of these analyses (Figs 4–8) is deposited as an integrity-hashed, self-contained study bundle that can be re-imported into *golgi* and reproduced byte-for-byte (Data availability).

**Fig 5.**
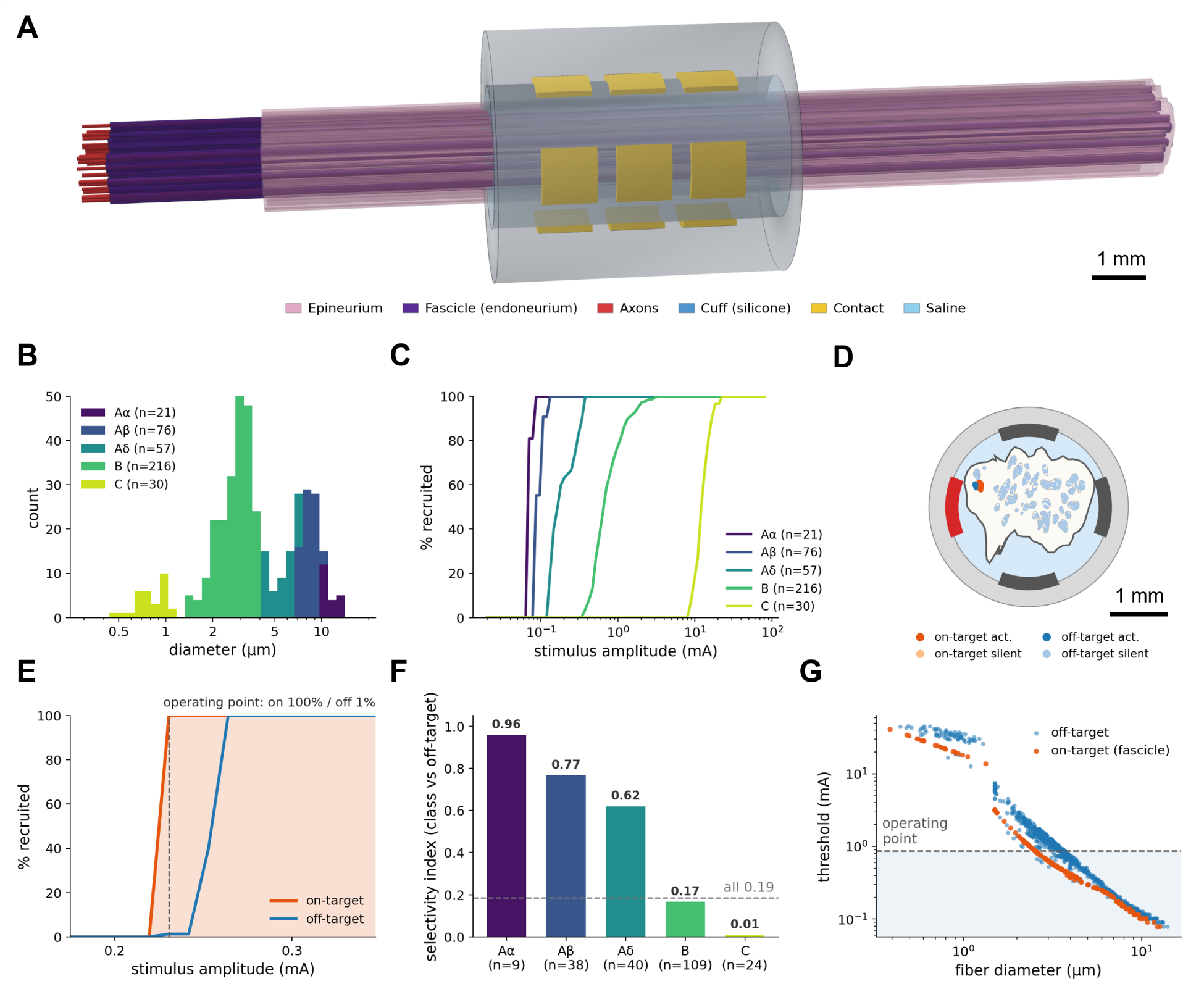
Image-derived swine cervical vagus: population recruitment and fascicular selectivity. **A:** Cel-shaded rendering of the extruded multifascicular nerve within the 12-contact cuff, with the sampled fiber population. **B:** Fiber-diameter distribution by axon class (A*α*/A*β*/A*δ*/B/C). **C:** Recruitment by fiber class versus stimulus amplitude (central contact, 1 ms cathodic monophasic pulse), reproducing the large-myelinated-first, C-fiber-last order. **D:** Fascicular-targeting cross-section (uniform 10 µm fibers)—fascicle boundaries with the peripheral target fascicle shaded (on-target) versus the remainder (off-target); fibers styled by on/off-target × activated/silent, optimal contact marked on the cuff. **E:** On-versus off-target recruitment versus amplitude (uniform 10 µm fibers), with the selective operating point marked (target ∼100% / off ∼1%). **F:** Per-class selectivity index (target fascicle versus off-target) for the realistic population: the large myelinated A*α* (0.96) and A*β* (0.77) fibers are selectively recruited, the small cardiac B (0.17) and unmyelinated C (0.01) fibers far less. **G:** Per-fiber activation threshold versus fiber diameter (realistic population), colored on-versus off-target, with the operating-point band; the selectively recruited fibers are the large, low-threshold ones. *Sample:* swine cervical vagus sub-4/sam-3, SPARC pig vagus morphology dataset [51].

**Fig 6.**
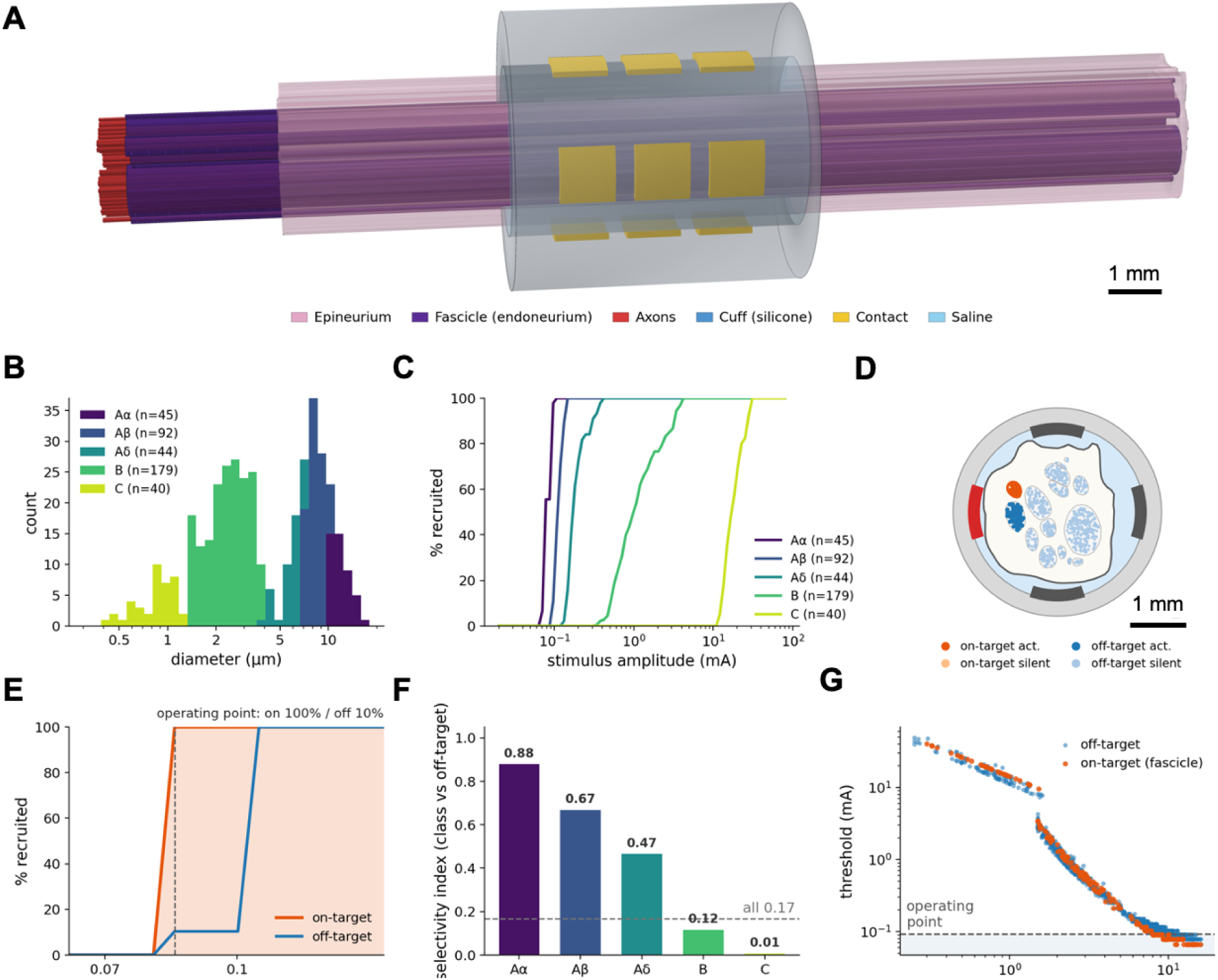
Image-derived human cervical vagus: population recruitment and fascicular selectivity. Panels as in Fig 5: **A:** cel-shaded render of the nerve in the 12-contact cuff; **B:** fiber-diameter distribution by axon class; **C:** recruitment by class; **D:** fascicular-targeting cross-section (uniform 10 µm fibers; target shaded, optimal contact marked); **E:** on-versus off-target recruitment versus amplitude (uniform 10 µm fibers, operating point marked); **F:** per-class selectivity index for the realistic population (large myelinated A*α* 0.88, A*β* 0.67 selectively recruited; small cardiac B 0.12, C 0.01 far less); **g** per-fiber threshold versus diameter for the realistic population—for the human cervical vagus nerve. *Sample:* human cervical vagus sub-50/sam-2, SPARC human vagus morphology dataset [52].

**Fig 7.**
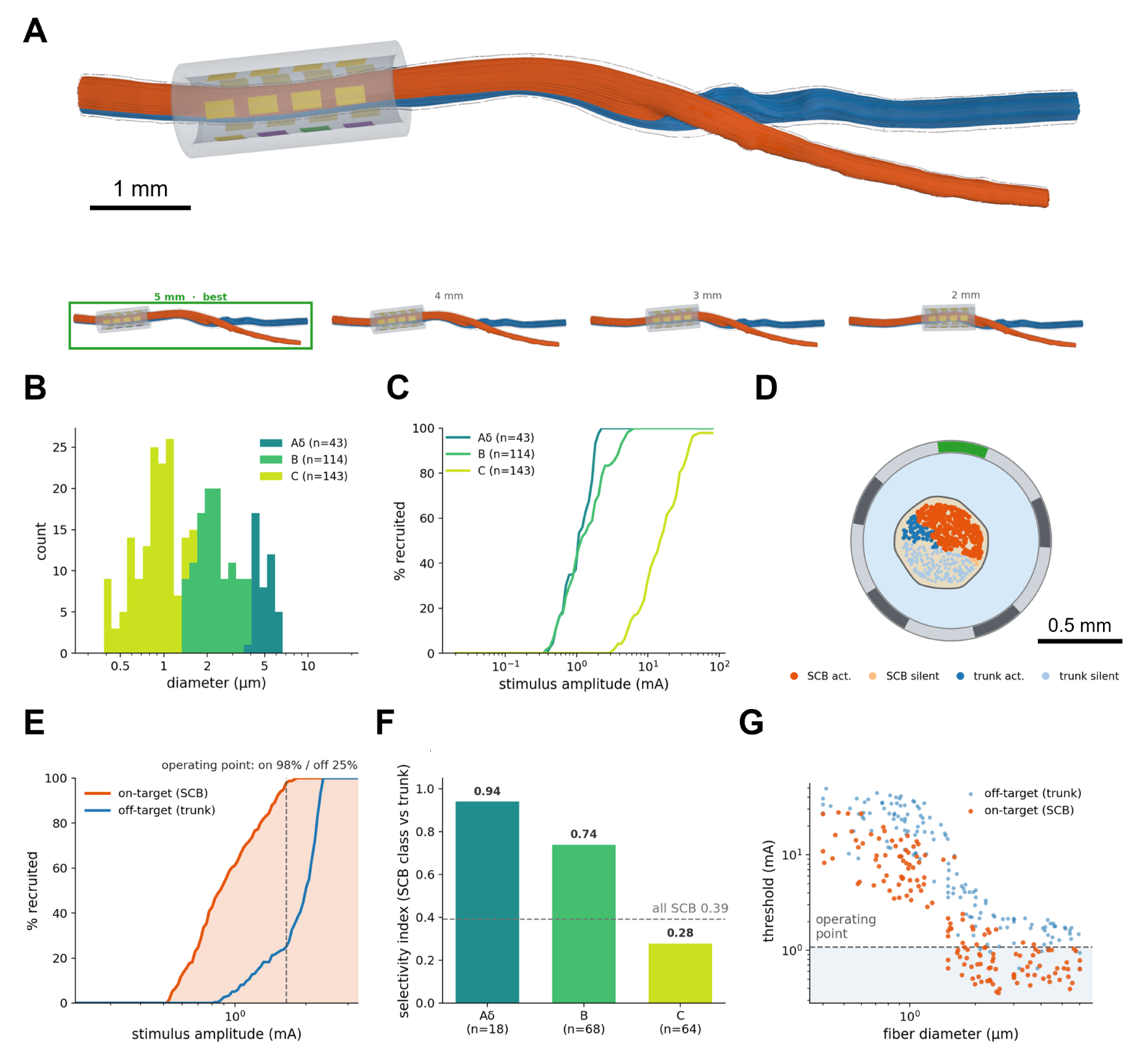
Branch-selective stimulation in a real three-dimensional rabbit nerve. **A:** Micro-CT-reconstructed rabbit cervical vagus with the 20-contact (4×5) ring-array cuff on the common trunk and streamline-traced fibers coloured by destination branch (orange, superior cardiac branch [SCB]; blue, trunk continuation), which diverge at the distal bifurcation; small panels below, the cuff swept to four longitudinal positions (labelled by distance from the bifurcation; the most selective, 5 mm, outlined). **B:** Fiber-diameter distribution by axon class for the rabbit-appropriate, C-dominated population (A*δ*/B/C). **C:** Recruitment by fiber class versus amplitude, reproducing the large-to-small order with C-fibers last. **D:** Cuff cross-section (uniform 10 µm fibers)—nerve boundary with fibers coloured by destination branch (orange SCB, blue trunk) and by activation state at the operating point, the tripole cathode (green) and its axial guard anodes marked; SCB- and trunk-bound fibers are partially segregated. **E:** On-target (SCB) versus off-target (trunk) recruitment versus amplitude (uniform 10 µm fibers) under the longitudinal tripole, with the selective operating point marked. **F:** Per-class SCB-to-trunk selectivity index (Veraart) for the realistic population under the longitudinal tripole: the small-myelinated A*δ* (0.94) and cardiac B (0.74) fibers are selectively recruited, the unmyelinated C-fibers least (0.28). **G:** Per-fiber activation threshold versus fiber diameter under the tripole, coloured on-target (SCB) versus off-target (trunk); the shaded band marks recruitment at the selective operating point. The SCB fibers (including small-diameter cardiac fibers) are recruited at lower currents than trunk fibers of the same size. *Sample:* rabbit cervical vagus micro-CT dataset [53].

**Fig 8.**
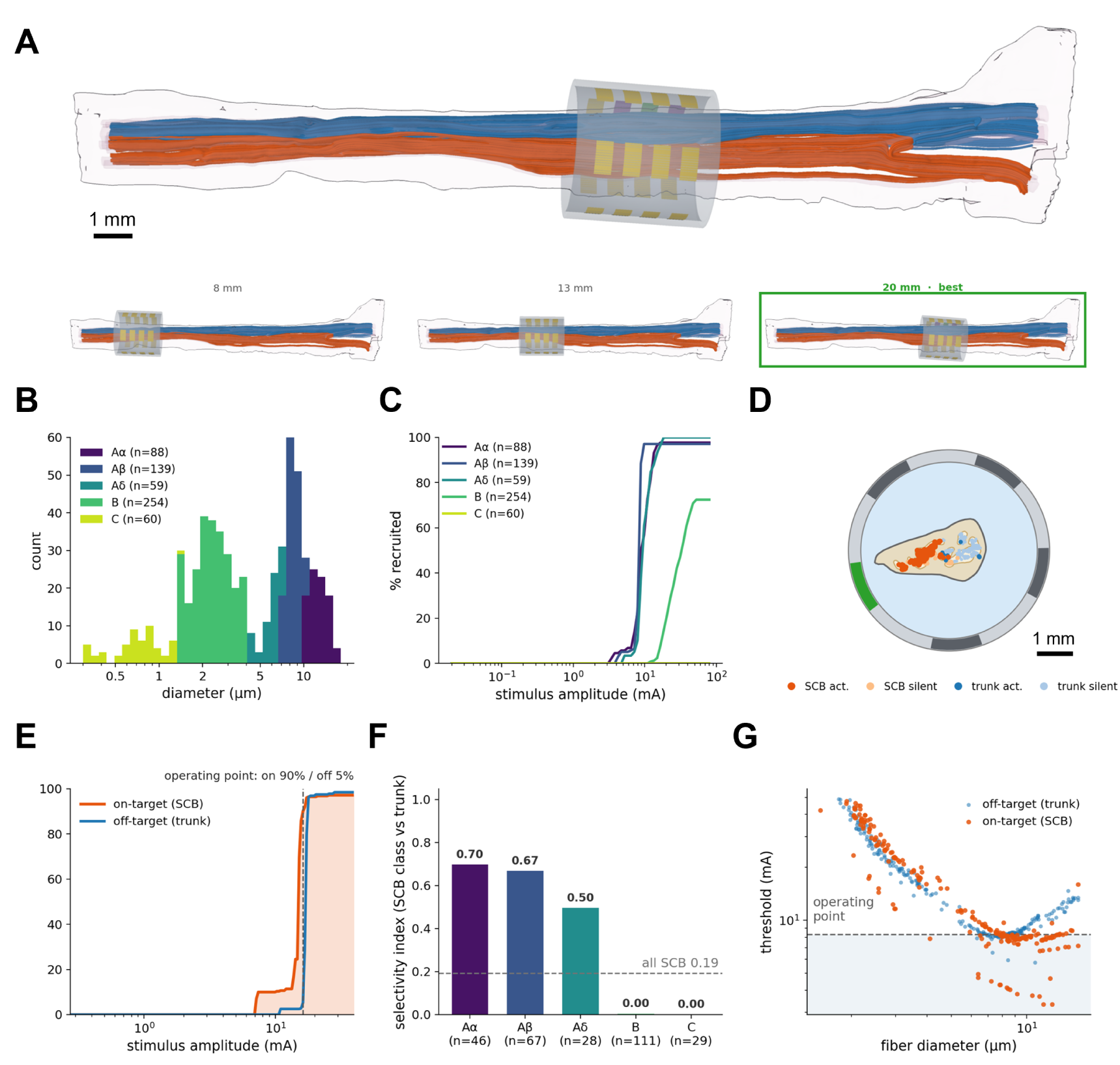
Branch-selective stimulation in a real three-dimensional human cervical vagus. **A:** Reconstructed human cervical vagus—segmented epineurium (translucent) enclosing the fascicular tree—with the 20-contact ring-array cuff on the proximal trunk and streamline-traced fibers coloured by destination branch (orange, superior cardiac branch [SCB]; blue, trunk continuation); small panels below, the cuff swept to three longitudinal positions (labelled by distance from the bifurcation; the most selective, 20 mm, outlined). **B:** Fiber-diameter distribution by axon class (A*α*/A*β*/A*δ*/B/C). **C:** Recruitment by fiber class versus amplitude (large-to-small order, C-fibers last). **D:** Cuff cross-section (uniform 10 µm fibers)—the multi-lobe fascicles with fibers coloured by destination branch (orange SCB, blue trunk) and activation state at the operating point; tripole cathode (green) and guard anodes marked. **E:** On-target (SCB) versus off-target (trunk) recruitment versus amplitude (uniform 10 µm fibers) under the longitudinal tripole, with the selective operating point marked. **F:** Per-class SCB-to-trunk selectivity index (Veraart) for the realistic population under the longitudinal tripole: only the large myelinated A*α* (0.70), A*β* (0.67) and A*δ* (0.50) fibers retain partial separability, whereas the small cardiac B (0.00) and unmyelinated C (0.00) fibers—whose high thresholds require currents that also recruit off-target fibers—are not separable. **G:** Per-fiber activation threshold versus fiber diameter under the tripole, coloured on-target (SCB) versus off-target (trunk); the shaded band marks recruitment at the selective operating point. Only the largest, lowest-threshold fibers are recruited, and the on- and off-target clouds overlap, reflecting the limited selectivity. *Sample:* human cervical vagus micro-CT dataset [54].

### From a multifascicular nerve image to a solved field

From a segmented multifascicular nerve image, *golgi* reconstructs the nerve within a multi-contact cuff, generates a multi-region finite-element mesh (Fig 3A), and solves the anisotropic extracellular field with perineurium contact impedance. The per-contact lead-field potential and electric field concentrate at the active contact and decay into the nerve (Fig 3B,C); sampling the potential along each fiber and twice differentiating along its arc length yields the activating function that localizes the drive to the region under the contact (Fig 3D). This establishes the field inputs reused, by superposition, for all subsequent fiber simulations. The identical pipeline was applied across a cohort of 36 nerve samples (18 swine, 18 human; 1–63 fascicles per nerve); per-sample fascicular anatomy and finite-element fields are provided in S2 Fig–S7 Fig and summarized in S1 Table; the two micro-CT-reconstructed nerves of this study are shown in the same style in S14 Fig.

### Realistic three-dimensional fiber trajectories

*golgi* generated fiber populations and trajectories spanning idealized and realistic anatomy (Fig 2B). For extruded nerves, fibers run straight (0 mm lateral deviation). For genuine three-dimensional samples—the human and rabbit cervical vagus nerves reconstructed from micro-CT imaging (see Data availability)—the quasi-static streamline solver produced smoothly curved trajectories that follow the endoneurial course—a median lateral deviation from the straight chord of 0.6 mm (rabbit) and 0.7 mm (human), up to 1.5 mm—and traversed the bifurcation (Fig 7A, 8A), which straight-fiber tools cannot represent. This streamline trajectory and branch-detection step is numerically converged (S11 Fig). Across fiber-seed counts from 100 to 1000 (three random seedings each), the median lateral deviation of the traced fibers was invariant (0.55 mm rabbit, 0.62 mm human) and the fraction of fibers assigned to the cardiac branch was stable (rabbit 0.46, human 0.42), with the realization spread narrowing from ±0.03 at 100 seeds to ±0.01 at 600. The trajectory geometry was likewise insensitive to the streamline integration step over 50–400 µm, and the per-class selectivity indices changed by less than 0.05 under two- to six-fold changes in fiber count, preserving every qualitative selectivity conclusion down to 50 fibers; branch detection requires only that the cap-clustering threshold be set below the inter-branch separation.

### Validation against analytic and experimental benchmarks

*golgi* reuses the field-standard governing equation, tissue conductivities, and biophysical fiber models established by prior peripheral nerve modeling work, so these components are validated by construction; the principal new engineering—an open FEniCSx finite-element solver in place of a commercial one—we validated directly. *golgi* solves the quasi-static Laplace equation [28] with literature tissue conductivities [30] (Table 1) and established axon models (MRG [11]; Sundt [14] and Tigerholm [15]). We cross-validated this solver directly against the commercial reference (COMSOL) on three independently rebuilt, re-meshed models (Fig 4A,B). For a current monopole in a grounded saline cylinder, *golgi* and COMSOL both reproduce the closed-form potential (near-field mean error 0.5 % and 1.3 %, respectively; Fig 4A). For an idealized cuff under full anisotropic, perineurium-contact-impedance physics and for a real swine cervical vagus, the *golgi* and COMSOL per-contact lead fields agree point-for-point to a mean of 2.6 % and 1.9 % (*R*^2^ ≥ 0.98; Fig 4B), confirming that the open FEniCSx stack—including the anisotropic conductivity and the perineurium contact-impedance boundary—reproduces the commercial solver (full per-model detail in S15 Fig). We separately confirmed the fiber-model foundations with two checks (S8 Fig): the conduction velocity of *golgi*’s interpolated MRG fibers reproduces the McIntyre 2002 discrete MRG model [11] to within 3 % across all eight standard diameters (5.7–16 µm), benchmarked through the identical extracellular pipeline (overall 5.1 m s*^−^*^1^ µm*^−^*^1^); the diameter-dependent ratio rises sub-linearly from ∼4 m s*^−^*^1^ µm*^−^*^1^ at small caliber to ∼5 at large, remaining below the single-ratio Hursh upper bound [42] (∼6 m s*^−^*^1^ µm*^−^*^1^); and the strength–duration relationship follows the Weiss–Lapicque law [43, 44] with a rheobase of 48 µA and a chronaxie of 151 µs, within the reported range for myelinated peripheral fibers.

We then validated the complete image-to-mesh-to-FEM-to-recruitment pipeline against three independent recruitment benchmarks that span electrode types, fiber models, and species (Fig 4). First, we reproduced the *in vivo* dog cervical VNS benchmark of Yoo et al. [45]—the same experiment used to validate ASCENT—directly from ASCENT’s published segmentation of the dog cervical vagus: we extruded their nerve and fascicle masks through *golgi*’s image-to-mesh-to-FEM pipeline (the same pipeline applied to the micro-CT nerves above), yielding a 1.7 mm fascicle within a 3.27 mm nerve with a perineurium contact-impedance sheet, a LivaNova-style separated bipolar cuff at the clinical 8 mm electrode separation, and a surrounding muscle volume, driven by a 300 µs monophasic cathodic pulse. Comparing *golgi*’s centroid-fiber thresholds against both the *in vivo* ENG data (mean ± SD) and ASCENT’s own COMSOL-based predictions for the same preparation (Fig 4C), *golgi* reproduces the clinically important large-myelinated-first, C-fiber-spared recruitment order (A-fiber 0.28 mA *<* fast-B 0.71 mA *<* slow-B 1.61 mA *<* C-fiber 25 mA): the A-fiber threshold lies within the in-vivo SD band (as does ASCENT’s) and the C-fiber at the upper edge of its broad in-vivo range (just above ASCENT), whereas the two B-fiber thresholds are underestimated roughly two-fold relative to both the in-vivo data and ASCENT. The two B-fibers (2.1 and 3.6 µm) lie below the MRG model’s nominal calibration range (< 5.7 µm) [11], where absolute thresholds are most sensitive to the fiber-model implementation—the interpolated MRG geometry and the NEURON-versus-PyFibers numerics—whereas the recruitment order that underlies fiber-type-selective stimulation is robust to these choices.

Second, we reproduced the intrafascicular benchmark used to validate NRV [9]: recruitment of myelinated fibers by a longitudinal intrafascicular electrode (LIFE) in a cat-tibial-sized monofascicular nerve, compared against the *in vivo* data of Nannini and Horch [46] and Yoshida and Horch [47] (Fig 4D). *golgi*’s native reciprocity finite-element solver computes the LIFE lead field directly—a 25 µm intrafascicular wire embedded in the anisotropic endoneurium—and the resulting MRG recruitment-versus-current curves at 20 and 50 µs fall within the in-vivo and NRV bands (50 % recruitment at 53 µA for 50 µs and 90 µA for 20 µs); the strength–duration recruitment-rate ratio from 50 to 20 µs is 2.0, matching the *in vivo* value (2.1) and NRV (2.4).

Third, we reproduced the multifascicular-cuff benchmark of Bucksot et al. [48]: per-fascicle recruitment in a five-fascicle nerve under a circumferential bipolar cuff (Fig 4E,F). Running Bucksot’s segmentation through the same *golgi* pipeline, *golgi* recovers per-fascicle activation thresholds in the published 0.1–1.5 mA range with the correct large-fiber-first diameter dependence, and reproduces the study’s central finding that a circumferential contact recruits the nerve near-uniformly: rotating the 270 *^◦^* contact by 180 *^◦^* (“inverted”) leaves the whole-nerve recruitment essentially unchanged while individual fascicles shift only modestly with their distance from the active conductor.

### Population recruitment and fascicular selectivity in the swine and human cervical vagus

We sampled realistic fiber populations (mixed A-, B-, and C-type fibers, with fiber-diameter distributions and the relative proportions of A-, B-, and C-fibers drawn from quantitative morphometry of the human and pig cervical vagus nerve [7, 49]) for the image-derived swine and human nerves (example nerves drawn from the 36-sample cohort; S2 Fig–S7 Fig) and characterized, for each, both population recruitment and the achievable fascicular selectivity (swine, Fig 5; human, Fig 6). Single-contact recruitment with a 1 ms cathodic (cathode-leading) monophasic pulse followed the expected inverse dependence on diameter—large myelinated fibers activated at the lowest amplitudes (∼0.1 mA), small myelinated fibers at intermediate amplitudes (∼0.4–2 mA), and unmyelinated C-fibers only at high amplitudes (∼11–25 mA)—reproducing the physiological large-to-small recruitment order with C-fibers recruited last (Fig 5B,C; Fig 6B,C), the basis for fiber-type-selective stimulation. Using the 12-contact cuff and per-contact lead fields, *golgi* then quantified how selectively individual fascicles can be targeted. For uniform 10 µm fibers—isolating the geometric selectivity—current steering to a chosen peripheral target fascicle recruited essentially all on-target fibers while sparing off-target fibers (on-target 100%, off-target ∼0–1% at the operating point; Fig 5D,E; Fig 6D,E), and more than 70% of fascicles (swine 71%, human 87%) were selectively targetable above a Veraart selectivity index [50] of 0.5. Resolving the realistic fiber population by class, however, only the large myelinated fibers retained this separability (swine A*α* 0.88, A*β* 0.67; human A*α* 0.99, A*β* 0.73), whereas the small cardiac B and unmyelinated C fibers—which require higher currents that also recruit off-target fibers—did not (swine B 0.17, C 0.01; human B 0.12, C 0.01; Fig 5F; Fig 6F); plotting each fiber’s threshold against its diameter shows this size dependence directly (Fig 5G; Fig 6G).

### Branch-selective stimulation of a real three-dimensional nerve

*golgi* resolves anatomically defined branches in a real three-dimensional rabbit cervical vagus nerve and poses a question that straight-fiber tools cannot: with a cuff on the common trunk, how selectively can the superior cardiac branch (SCB) be recruited while sparing the trunk continuation? The quasi-static streamline solver traced fibers from the trunk through the distal bifurcation into the two branches (232 to the SCB, 236 to the trunk continuation; Fig 7A) over a rabbit-appropriate fiber population dominated by small myelinated (A*δ*/B) and unmyelinated (C) fibers (Fig 7B), as is characteristic of the rabbit cervical vagus and consistent with our ongoing immunohistochemical characterization. This population recruited in the expected large-to-small order with C-fibers last (Fig 7C). To resolve per-class statistics we sampled the rarer small-myelinated cardiac classes more densely than their natural scarcity; this does not bias the per-class selectivity indices reported below, each of which is governed by the within-class threshold–diameter relationship rather than the inter-class proportions and is stable under two- to six-fold changes in fiber sampling (S11 Fig). Using a 20-contact (4×5) ring-array cuff and per-contact lead fields, *golgi* swept the cuff longitudinally along the common trunk and, at each position, compared monopolar and multipolar current-steering configurations (Fig 7A). A longitudinal guarded tripole—the SCB-coupled contact as cathode flanked by two axial guard anodes—gave the strongest, most robust steering. Selectivity was highest about 5 mm upstream of the bifurcation, where the SCB- and trunk-bound fibers are most angularly segregated within the trunk cross-section (Fig 7D), and declined as the cuff approached the branch (Veraart selectivity index [50] 0.73 falling to 0.39 from 5 to 2 mm; controlled 10 µm fibers, SCB 98% vs trunk 25% at the operating point; Fig 7D,E, S9 Fig). For the realistic mixed population the same tripole selectively recruited the clinically relevant small-myelinated cardiac fibers: the A*δ* and B (cardiac) SCB fibers reached per-class selectivity indices of 0.94 and 0.74—near-complete on-target recruitment with the trunk largely spared—whereas the unmyelinated C-fibers—the highest-threshold class and the least branch-segregated across the cross-section—were the least separable (0.28; Fig 7F). Plotting each fiber’s activation threshold against its diameter under the tripole shows the basis of this branch selectivity: across the diameter range the SCB (on-target) fibers—including the small-diameter cardiac fibers—are recruited at lower currents than the trunk fibers, so the operating point captures them while largely sparing the trunk (Fig 7G). This quantitative, anatomy-specific, branch-resolved analysis—steering from a proximal cuff to a named distal branch and resolving which fiber classes can be captured—is one that straight-fiber modeling tools cannot perform.

### Branch-selective stimulation of the right human cervical vagus in three dimensions

The same branch-resolved workflow, applied to a real three-dimensional human cervical vagus—reconstructed as a multi-region mesh in which the segmented epineurium encloses the fascicular tree (Fig 8A,D)—targeted the SCB from a 20-contact ring-array cuff swept along the proximal trunk. *golgi* traced full-length fibers from the trunk into the SCB and trunk continuation (281 SCB, 319 trunk; Fig 8A) and sampled a human vagal population spanning A*α* through C fibers, with diameter spectrum and superior-cardiac-branch fiber organization following the morphometry of Kronsteiner et al. [49] (Fig 8B,C). At the proximal trunk the SCB- and trunk-bound fibers lie close together, so a single contact, whose field is not spatially confined, recruited only a small fraction of the SCB selectively. For uniform 10 µm fibers—isolating the geometric targeting—a longitudinal guarded tripole steered the extracellular field to the SCB cluster and recruited it while largely sparing the trunk continuation (SCB 90% vs trunk 5% at the operating point; Veraart selectivity index 0.85; Fig 8D,E; S10 Fig). For the realistic population, however, only the large myelinated fibers retained that separability (A*α* 0.70, A*β* 0.67, A*δ* 0.50), whereas the small cardiac B and unmyelinated C fibers—whose high thresholds require currents that, at the levels needed to recruit them, also recruit the off-target large myelinated fibers throughout the trunk—were not separable at all (B 0.00, C 0.00; Fig 8F); plotting each fiber’s threshold against its diameter shows this size dependence directly (Fig 8G). This contrasts with the rabbit, whose cardiac fibers are predominantly small and whose SCB is a separate branch, so that even the small cardiac B-fibers remain selectable. *golgi* thus delivers quantitative, fiber-class-resolved branch-selective targeting on a fully reconstructed three-dimensional anatomy of the nerve most relevant to clinical vagus nerve stimulation.

To quantify what the full three-dimensional reconstruction adds over the conventional extruded-slice approximation [38], we built an extruded counterpart of the same human nerve: the cross-section under the depolarizing (cathodic) tripole contact—epineurium and five fascicles—extruded into straight, constant-section tubes, then meshed and solved through the identical pipeline with the same 20-contact array, longitudinal tripole, perineurium contact impedance, and fiber population, and seeded with straight axons at the same cuff-plane positions carrying the same branch labels (S13 Fig). Holding everything but the trajectory geometry fixed, the extruded approximation reproduced the reconstruction’s per-class selectivity to within 0.03 for the A*β*, A*δ*, B, and C fibers and preserved every qualitative conclusion, but overestimated large-fiber (A*α*) selectivity by 0.15 (extruded 0.85 versus reconstructed 0.70). Because both models used identical branch labels—and the SCB fibers are in fact ∼95% separable by fascicle in this nerve, so labeling is not the difference—the gap reflects the straight-extrusion idealization itself: perfectly cuff-aligned axons sharpen the longitudinal-tripole activating function and hold off-target trunk A*α* fibers apart (off-target recruitment 9% versus 17% at the operating point), whereas the real curved, branching trajectories carry target and non-target large fibers together through the cuff. The full reconstruction therefore yields a more conservative, anatomically grounded large-fiber selectivity estimate, while confirming that an extruded slice suffices for the smaller, less excitable classes. This single-nerve comparison is consistent with the systematic extrusion-versus-true-3D analysis of human VNS by Marshall et al. [38]: as in their study, the slice under the cathodic contact—the choice they identify as essential for agreement—reproduced most of the true-3D response, and the residual large-fiber discrepancy falls squarely in the two regimes they report as least faithful to extrusion, namely a longitudinally distributed current-steering contact arrangement (our guarded tripole imposes the cathode-plane cross-section on guard anodes 1 mm away, where the human vagus splits or merges every ∼0.56 mm) evaluated on the acute, undeformed nerve rather than a chronically cuff-deformed circular cross-section.

### Verifiable, shareable study bundles

A complete swine study—mesh, field, fibers, and results—was exported as a single integrity-hashed bundle. The manifest captured the per-stage processing graph with a SHA-256 hash of every file. Verifying the intact bundle with golgi replay confirmed all files matched their recorded hashes; deliberately altering a single byte of one file caused replay to detect the mismatch and fail. To our knowledge no other open peripheral nerve modeling platform ships integrity-hashed, replayable study bundles; the bundle format and verification workflow are detailed in an accompanying software paper. The golgi study bundles behind the image-derived and micro-CT nerve simulations in this work (Figs 4–8)—each carrying the geometry, mesh, finite-element lead fields, fiber population, and recruitment behind the figure—are deposited on Zenodo [55] and can be imported into *golgi* and re-verified byte-for-byte, so each result can be reopened and reproduced from the exact inputs that generated it.

## Discussion

*golgi* addresses a practical gap in peripheral nerve stimulation modeling: existing open tools are realistic but code-first, and some depend on commercial solvers, which limits who can build and run nerve models [8, 9]. By exposing the entire image-to-recruitment pipeline through a graphical interface—while preserving a scriptable API and command-line interface for batch and high-performance use—*golgi* makes anatomically realistic modeling accessible to experimentalists and clinicians without sacrificing rigor. The demonstrations reproduce established physiology (the inverse diameter–threshold relationship and large-to-small recruitment order [10, 11]) and quantify fascicular selectivity and current steering with a multi-contact cuff [6, 50, 56], indicating that the accessibility gains do not come at the expense of model fidelity. Notably, NRV—the closest open peer, sharing *golgi*’s open Python, FEM, and NEURON backbone—uses idealized parametric fascicle geometry and a single CPU backend, and explicitly lists realistic image-derived geometry and GPU acceleration among its future directions [9]; *golgi* delivers both today.

Two capabilities distinguish *golgi* among open tools. First, *golgi* brings physically curved three-dimensional fiber trajectories and anatomical-branch detection into a fully open, no-code package. That true three-dimensional morphology—fascicle splitting and merging and curved fiber paths—alters human VNS predictions has very recently been established by Marshall et al. [38], whose pipeline likewise builds curved-3D models from micro-CT; it depends, however, on commercial meshing and finite-element solvers (Simpleware, COMSOL) and is operated from code, whereas *golgi* delivers the same curved-3D, branch-resolved capability through a fully open solver stack and a graphical interface. This enables branch-targeted analyses in genuinely three-dimensional, bifurcating nerves that straight-fiber pipelines cannot represent. Applied to vagal cardiac targeting, this yielded a concrete, anatomy-dependent result: a discrete branch such as the rabbit superior cardiac branch can be selectively recruited from a proximal cuff down to its small cardiac (B-type) fibers, whereas in the human cervical vagus—where isolating the small cardiac fibers would require currents that also recruit off-target fibers—current steering isolates only the large myelinated fibers. This distinction, which only a branch-resolved three-dimensional model can expose, has direct implications for where and how cardiac-selective vagus nerve stimulation is feasible [7, 57]. The converse of this cardiac result is equally useful: the same fiber-type-resolved, three-dimensional selectivity analysis is directly relevant to the indications for which left cervical VNS is already in clinical use—rheumatoid arthritis [3], epilepsy [1], and depression [2]—where left-sided placement itself reflects a need to spare cardiac fibers and side-effect-limited dosing constrains efficacy. There, *golgi* offers a per-anatomy route to tune contact configuration and current steering for the intended therapeutic recruitment while minimizing off-target activation. Second, *golgi* packages each study as an integrity-hashed, independently verifiable bundle, turning end-to-end reproducibility and FAIR data sharing [58] from an aspiration into a one-command check.

### Limitations and future directions

*golgi*’s quantitative predictions inherit the assumptions of the underlying fiber and tissue models, and systematic experimental validation against measured recruitment is ongoing. The perineurium contact impedance is applied consistently across both solver paths—the forward field solver (validation models) and the per-contact reciprocity lead fields used for the multifascicular fascicular- and branch-selectivity analyses (Figs 5, 6, 8)—as a single representative thin-layer sheet resistance per nerve rather than a per-fascicle value. The monofascicular rabbit model of Fig 7 does not include this perineurium sheet—its single fascicle boundary is modeled as a plain conductivity interface—whereas the human model does, raising the question of whether the rabbit-versus-human selectivity contrast reflects this difference in treatment rather than anatomy. We tested this directly by re-solving the human branch-selectivity analysis with the perineurium contact impedance removed, matching the rabbit’s treatment: removing it sharpened large-fiber selectivity (A*α* rising from 0.70 to 0.97) but left the small cardiac fibers non-separable (B from 0.00 to 0.09, C unchanged at 0.00; S12 Fig). The perineurium treatment thus shifts any per-class selectivity index by at most 0.28 (0.09 for B fibers)—far smaller than the rabbit-human B-fiber contrast (0.74 versus 0.00)—so the contrast is intrinsic to the two nerves—the rabbit’s discrete, small-fiber cardiac branch versus the human’s clustered branch within a full fiber-size spectrum—not an artifact of perineurium treatment. The reciprocal control, adding a perineurium sheet to the rabbit, was not feasible on the sub-millimeter, curved, branching rabbit geometry, which does not admit a cleanly resolved endoneurium–epineurium shell on the present mesher. The integrated segmentation workflow accelerates geometry creation but still benefits from expert review. Current backends cover validated NEURON models and a GPU surrogate [23]; a differentiable backend is planned. Future work will extend validation across species and electrode types and broaden the library of fiber models and electrode geometries.

## Data availability

The *golgi* source code is publicly available on GitHub (https://github.com/CellularSyntax/golgi), together with the scripts that reproduce all figures and studies in this work, and is documented in the repository wiki (https://github.com/CellularSyntax/golgi/wiki). The release associated with this paper (v1.0.0), along with other archived releases, is deposited on Zenodo [59]; the most recent commit—there may be several per release—is available on GitHub. *golgi* runs on Linux, macOS, and Windows and is released under the AGPL-3.0-or-later license.

The micro-computed tomography (micro-CT) imaging datasets of the human and rabbit cervical vagus nerves used to construct the realistic three-dimensional geometries are deposited on Zenodo (rabbit [53], human [54]), each comprising the micro-CT image stacks, tissue segmentation masks, and reconstructed three-dimensional surface models. The image-derived swine and human cervical vagus geometries used for the cohort analyses (Figs 5, 6; S2 Fig–S7 Fig; S1 Table) are taken from the publicly available SPARC *Quantified vagus nerve morphology across species* resource—the pig [51] and human [52] datasets of Pelot and colleagues, whose primary publication is Pelot et al. [7].

The complete, self-contained *golgi* study bundles underlying the nerve simulations in this work are deposited on Zenodo [55]: each is an integrity-hashed project—geometry, mesh, finite-element lead fields, fiber population, and recruitment—that can be imported directly into *golgi* (via *Import study* on the File menu) and re-verified byte-for-byte with golgi replay, so every reported result can be reopened and reproduced from the exact inputs that generated it. The data accompanying this work—the micro-CT datasets and the *golgi* study bundles, all deposited on Zenodo—are released under the Creative Commons Attribution 4.0 International (CC-BY-4.0) license; this is distinct from the *golgi* software license (AGPL-3.0-or-later).

## Author contributions

**Conceptualization:** M.H. **Methodology:** M.H. **Investigation:** D.L., Y.J., R.B., L.R., L.Z., P.H., C.K., A.M., M.F., M.H. **Formal analysis:** D.L., Y.J., M.H. **Data curation:** D.L., Y.J., M.H. **Funding acquisition:** M.H. **Supervision:** M.H. **Writing – original draft:** D.L., Y.J., R.B., L.R., L.Z., P.H., C.K., A.M., M.F., M.H. **Writing – review & editing:** D.L., Y.J., R.B., L.R., L.Z., P.H., C.K., A.M., M.F., M.H.

## Acknowledgments

Simulations were performed using the Medical University of Vienna high-performance computing cluster. Yuting Jia gratefully acknowledges the financial support from the China Scholarship Council (CSC), File No. 202408460012.

## Supporting information

**S1 Fig.**
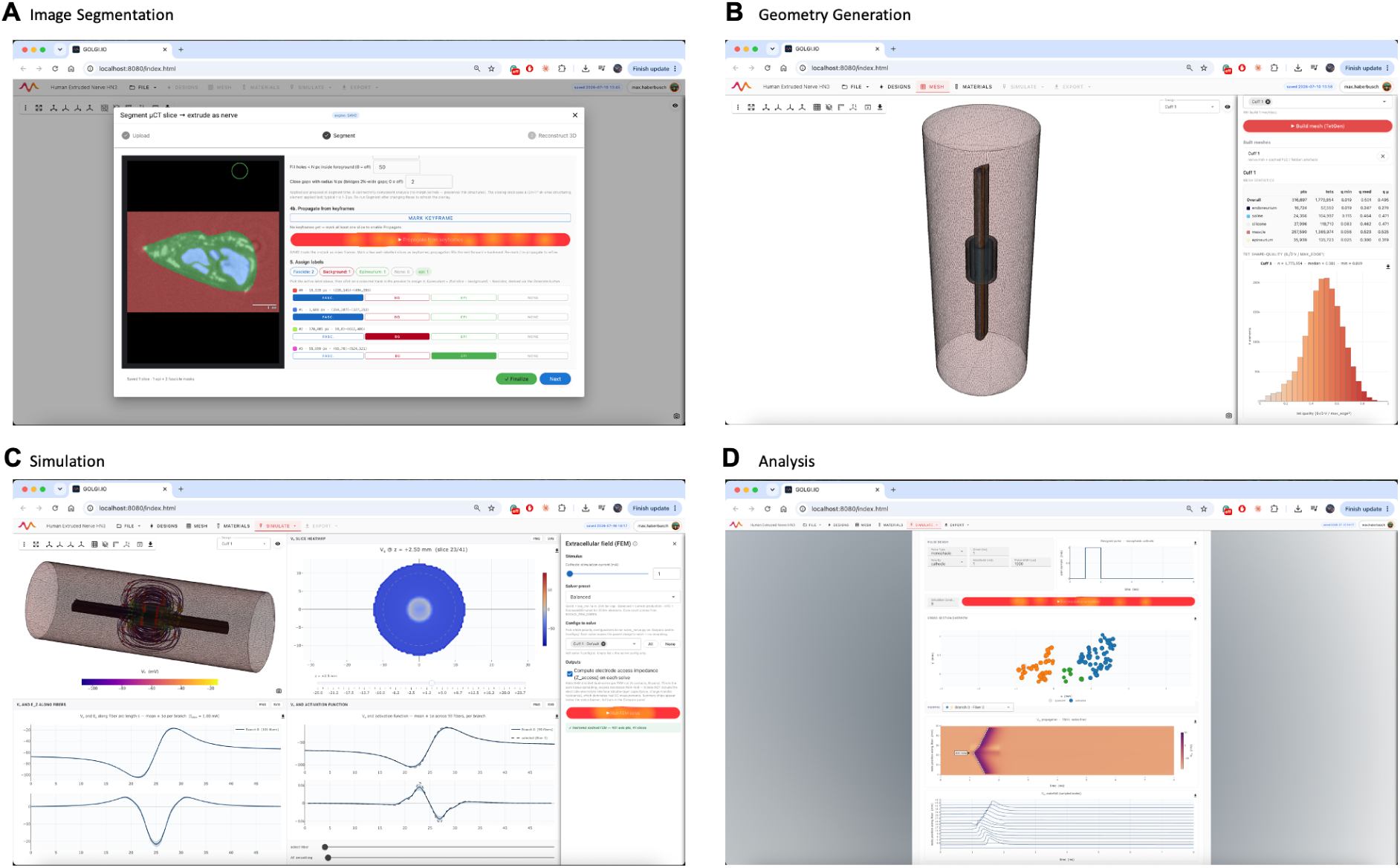
Graphical user interface. Representative views of the *golgi* graphical user interface across the four stages of the simulation workflow. **A:** Image segmentation, **B:** Multi-region geometry generation and meshing, **C:** Finite element simulation, and **D:** Analysis of the predicted nerve fiber activation.

**S2 Fig.**
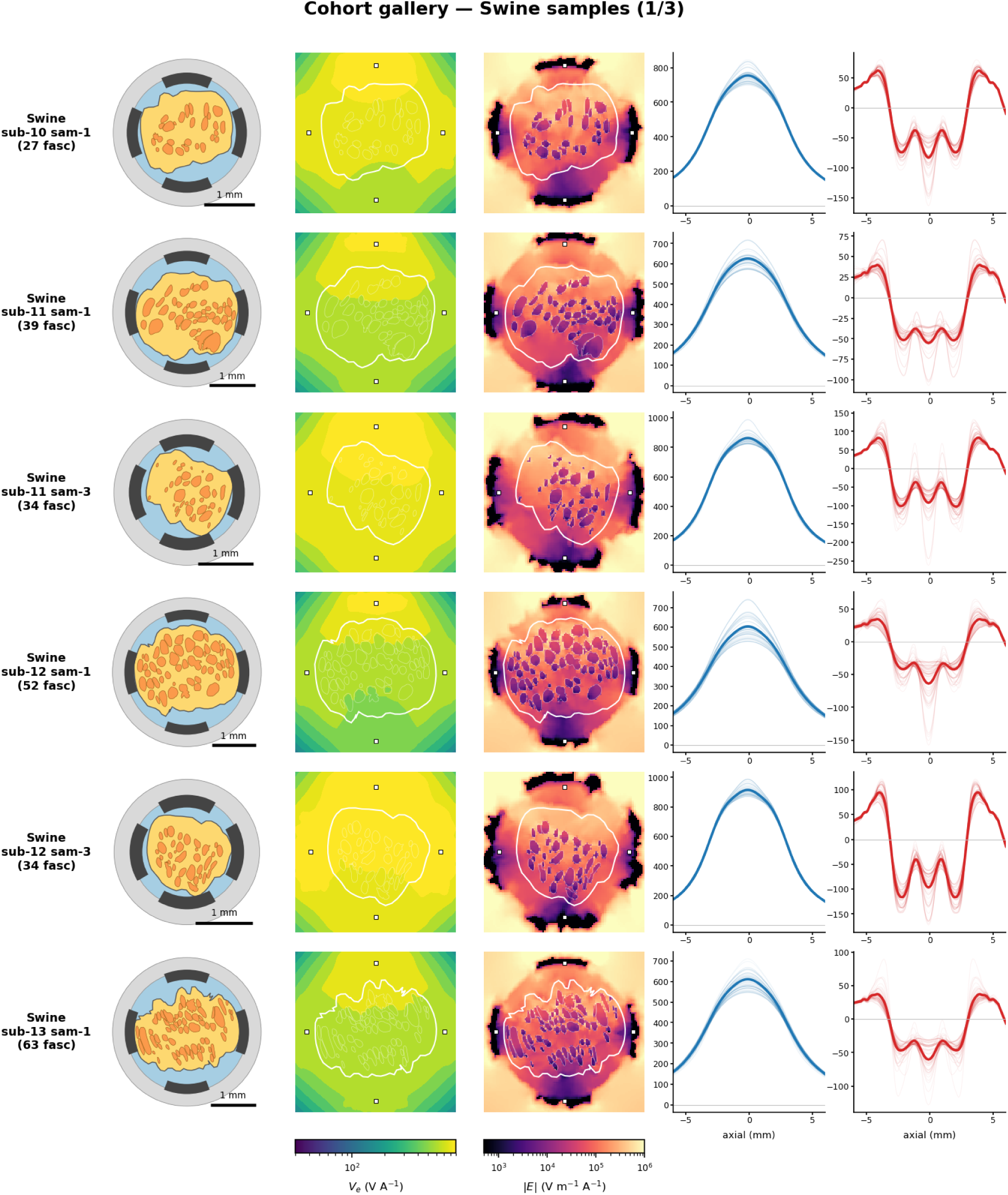
Cohort gallery, swine samples (1 of 3). For each sample: fascicle cross-section, finite-element lead-field *V_e_*, and electric-field magnitude |*E*| on the central cuff cross-section (*z* = 0).

**S3 Fig.**
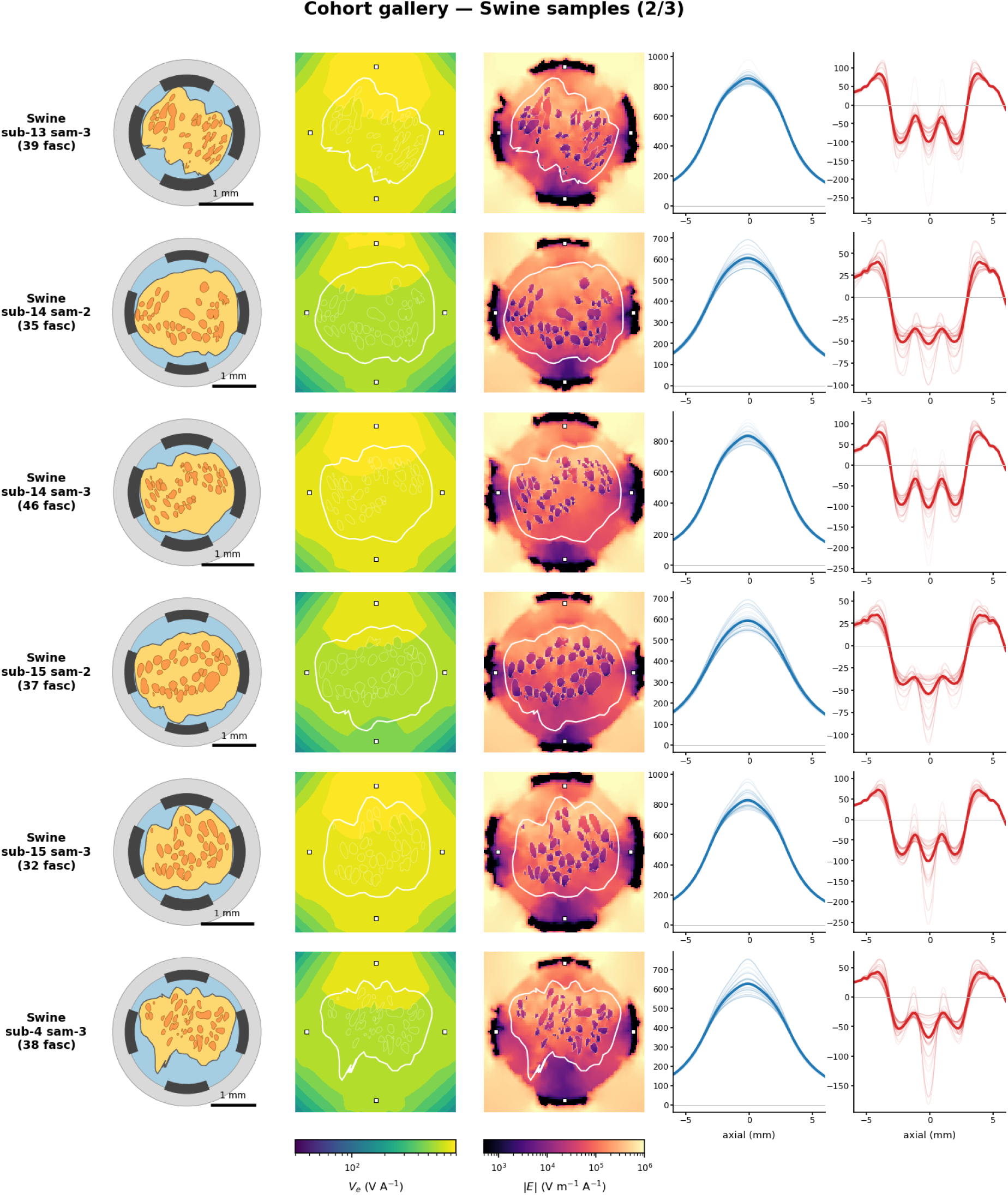
Cohort gallery, swine samples (2 of 3). For each sample: fascicle cross-section, finite-element lead-field *V_e_*, and electric-field magnitude |*E*| on the central cuff cross-section (*z* = 0).

**S4 Fig.**
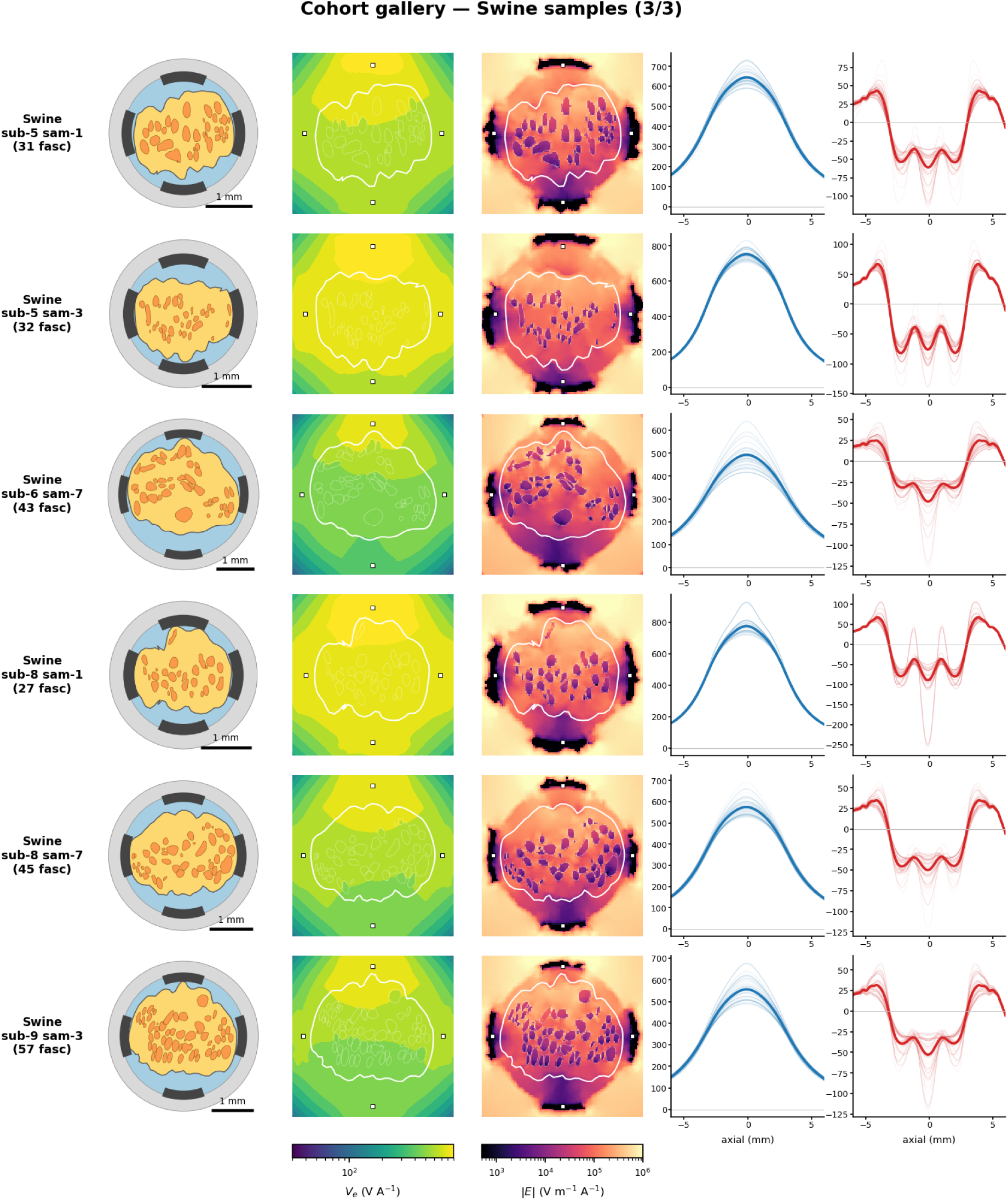
Cohort gallery, swine samples (3 of 3). For each sample: fascicle cross-section, finite-element lead-field *V_e_*, and electric-field magnitude |*E*| on the central cuff cross-section (*z* = 0).

**S5 Fig.**
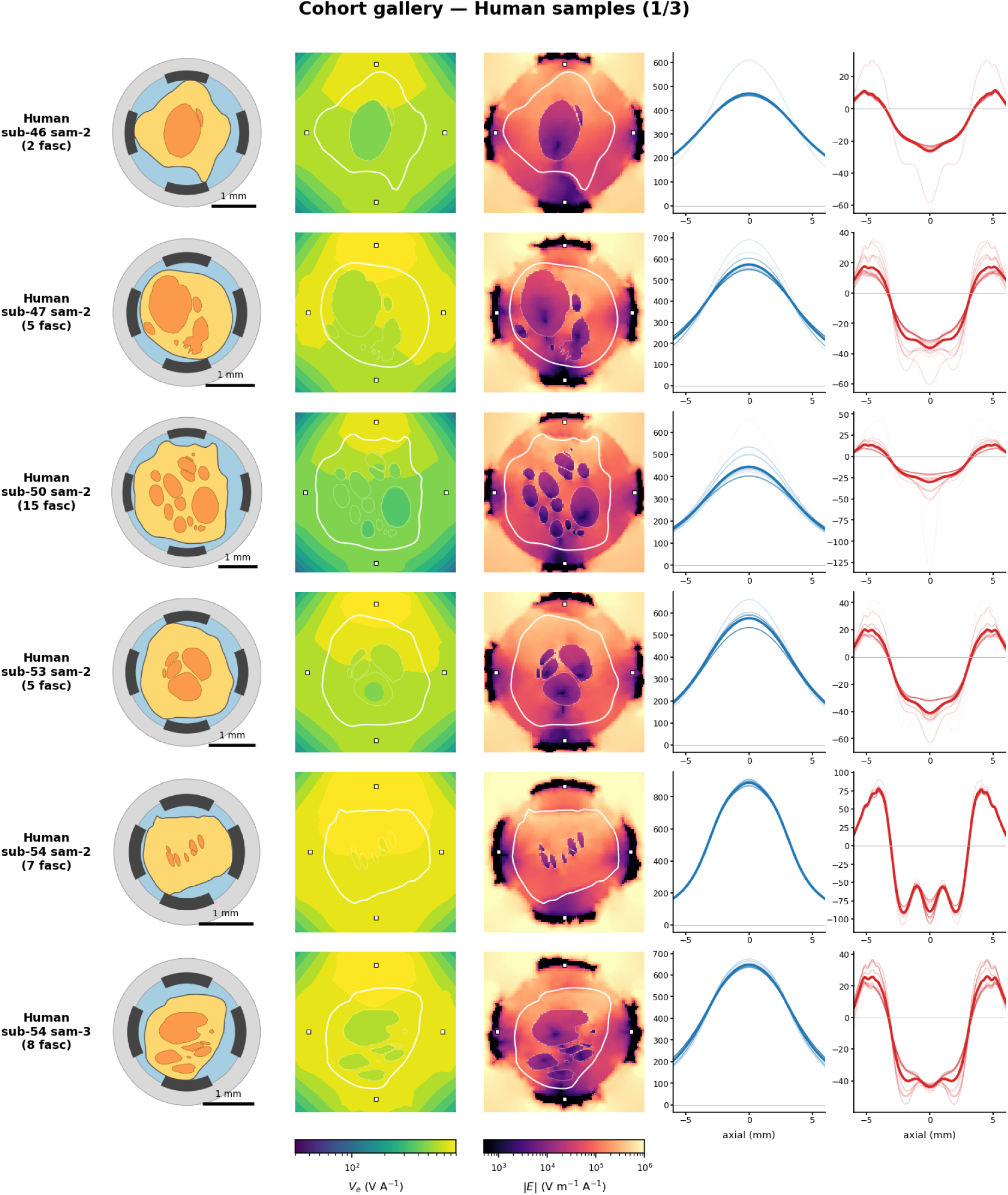
Cohort gallery, human samples (1 of 3). For each sample: fascicle cross-section, finite-element lead-field *V_e_*, and electric-field magnitude |*E*| on the central cuff cross-section (*z* = 0).

**S6 Fig.**
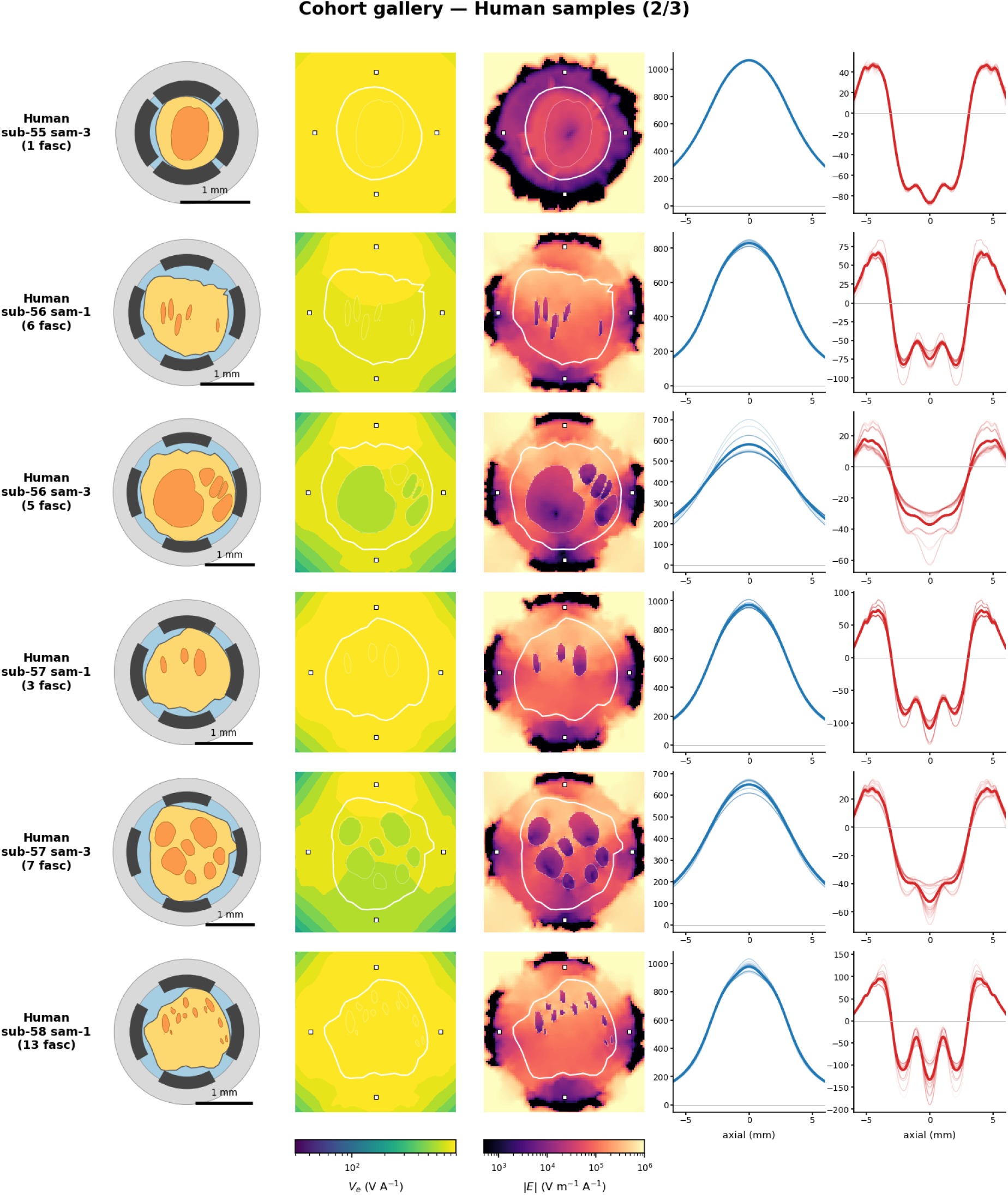
Cohort gallery, human samples (2 of 3). For each sample: fascicle cross-section, finite-element lead-field *V_e_*, and electric-field magnitude |*E*| on the central cuff cross-section (*z* = 0).

**S7 Fig.**
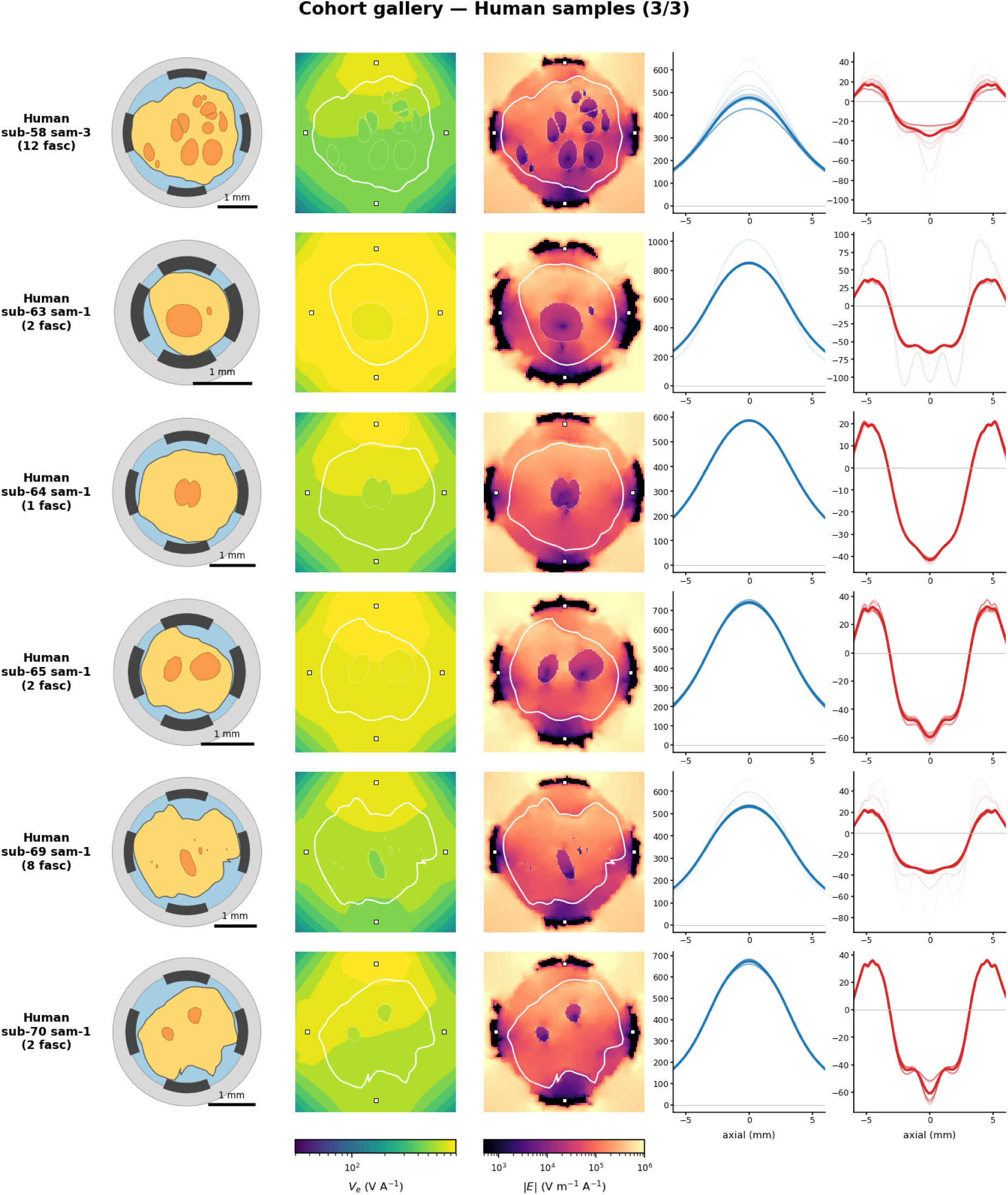
Cohort gallery, human samples (3 of 3). For each sample: fascicle cross-section, finite-element lead-field *V_e_*, and electric-field magnitude |*E*| on the central cuff cross-section (*z* = 0).

**S8 Fig.**
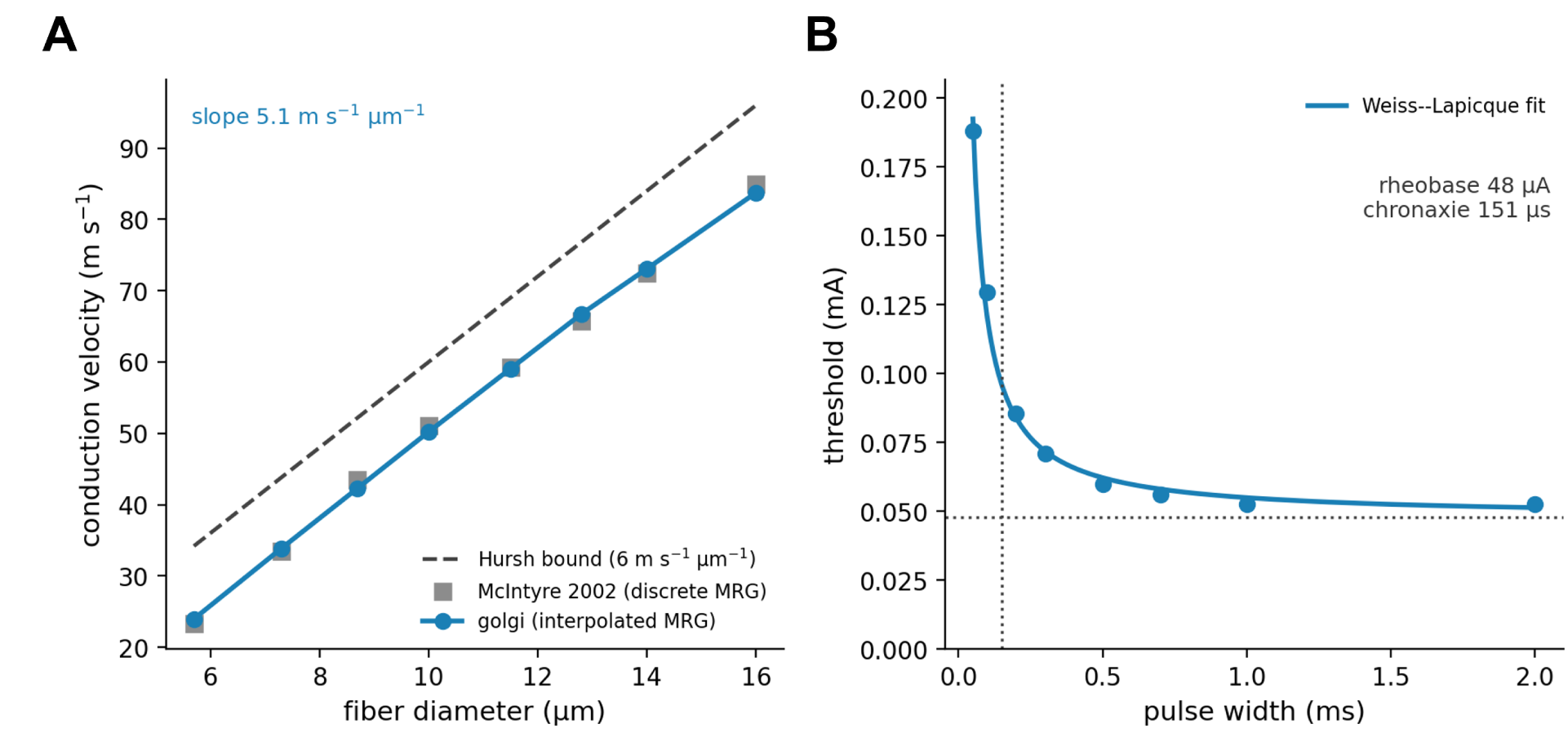
Fiber-model foundations. The two MRG fiber-model checks (the numerical solver accuracy is validated against the analytic monopole and COMSOL in Fig 4A,B and S15 Fig). **A:** Conduction velocity versus fiber diameter for *golgi* (interpolated PyFibers MRG) and the McIntyre 2002 MRG model (computed here from the discrete PyFibers MRG through the identical extracellular pipeline, so the two are directly comparable), with the Hursh rule shown for reference. *golgi* reproduces the discrete-MRG conduction velocities to within 3 % across all eight standard diameters (5.7–16 µm); the diameter-dependent ratio rises sub-linearly from ∼4 m s*^−^*^1^ µm*^−^*^1^ at small calibre to ∼5 at large, remaining below the single-ratio Hursh bound (6 m s*^−^*^1^ µm*^−^*^1^). **B:** Strength–duration curve for a 10 µm myelinated fiber with the Weiss–Lapicque fit (rheobase 48 µA, chronaxie 151 µs).

**S9 Fig.**
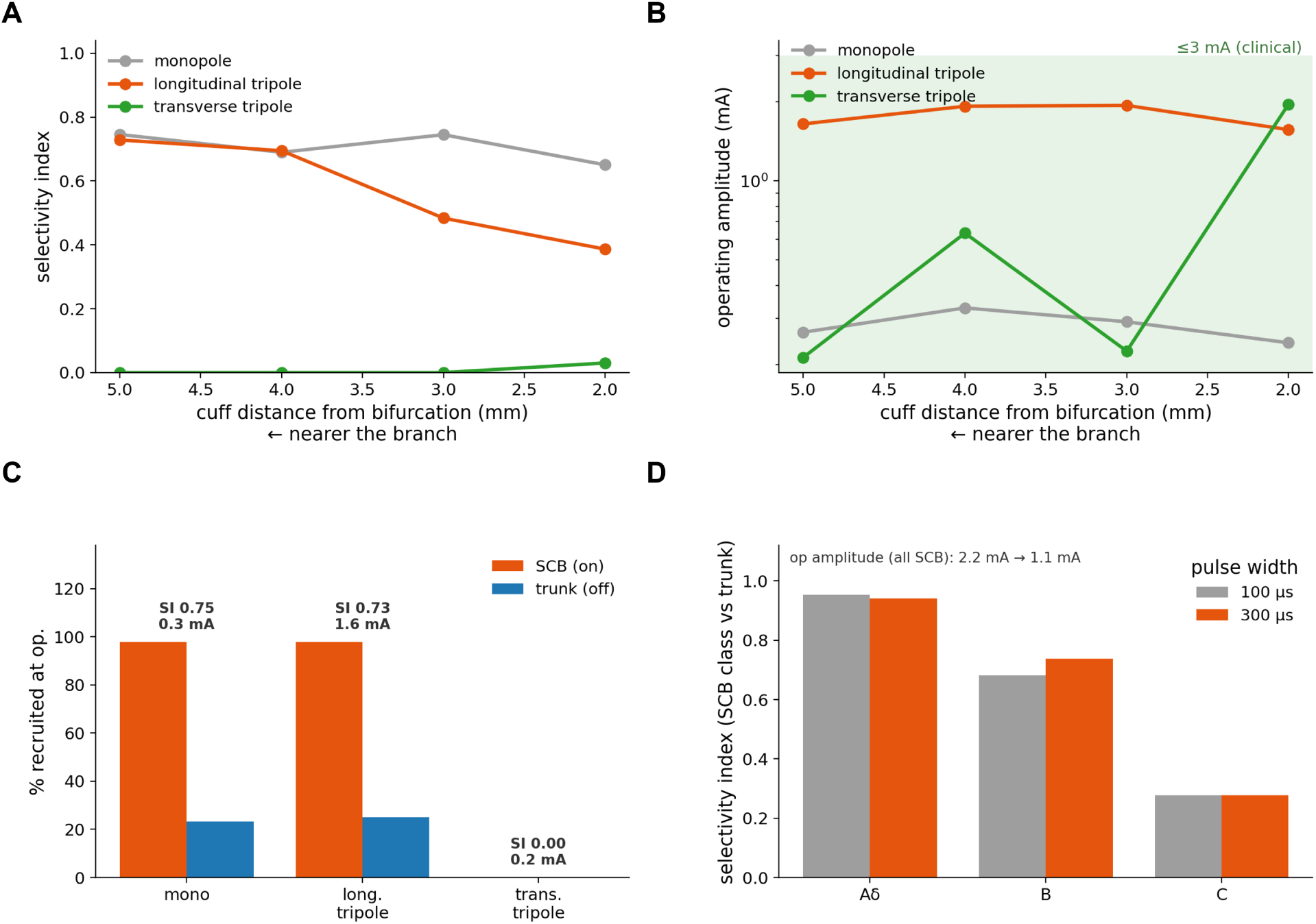
Rabbit superior-cardiac-branch selectivity: cuff-position × configuration sweep and pulse-width comparison. Controlled 10 µm fibers unless noted. **A:** Veraart selectivity index versus cuff distance from the bifurcation for monopolar, longitudinal-tripole, and transverse-tripole steering—selectivity is highest farthest upstream of the branch, and the transverse tripole is ineffective on the sub-millimetre rabbit cuff, where its three closely spaced angular contacts produce near-cancelling lead fields. **B:** Corresponding operating amplitude versus position (clinical ≤3 mA band shaded). **C:** On-versus off-target recruitment by configuration at the most selective position. **D:** Per-class selectivity index for the realistic population at 100 versus 300 µs pulse width—the longer (clinical) pulse lowers the operating amplitude while preserving small-myelinated (B) selectivity.

**S10 Fig.**
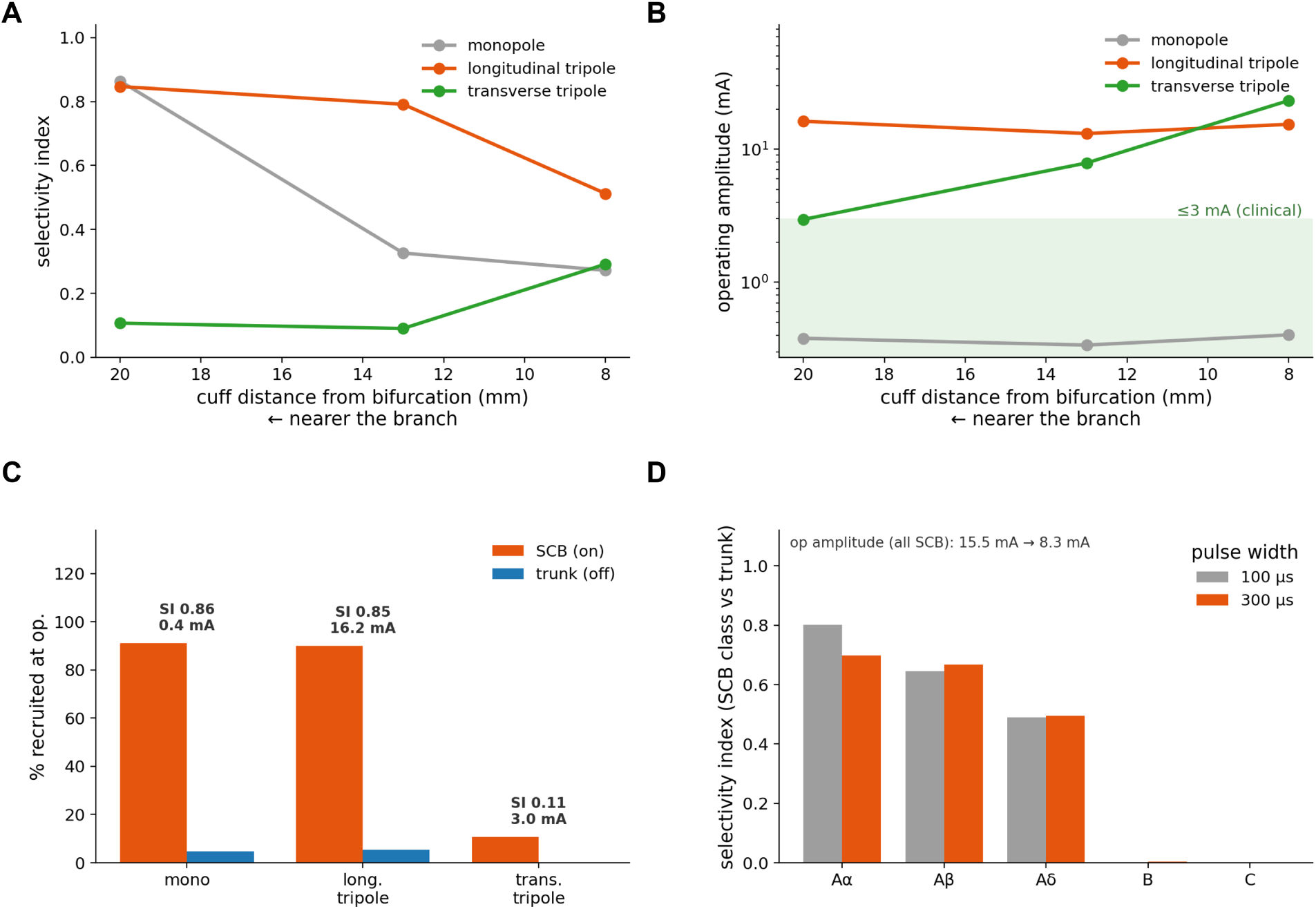
Human superior-cardiac-branch selectivity: cuff-position × configuration sweep and pulse-width comparison. Controlled 10 µm fibers unless noted. **A:** Veraart selectivity index versus cuff distance from the bifurcation for monopolar, longitudinal-tripole, and transverse-tripole steering. **B:** Corresponding operating amplitude versus position (clinical ≤3 mA band shaded). **C:** On-versus off-target recruitment by configuration at the most selective position. **D:** Per-class selectivity index for the realistic population at 100 versus 300 µs pulse width.

**S11 Fig.**
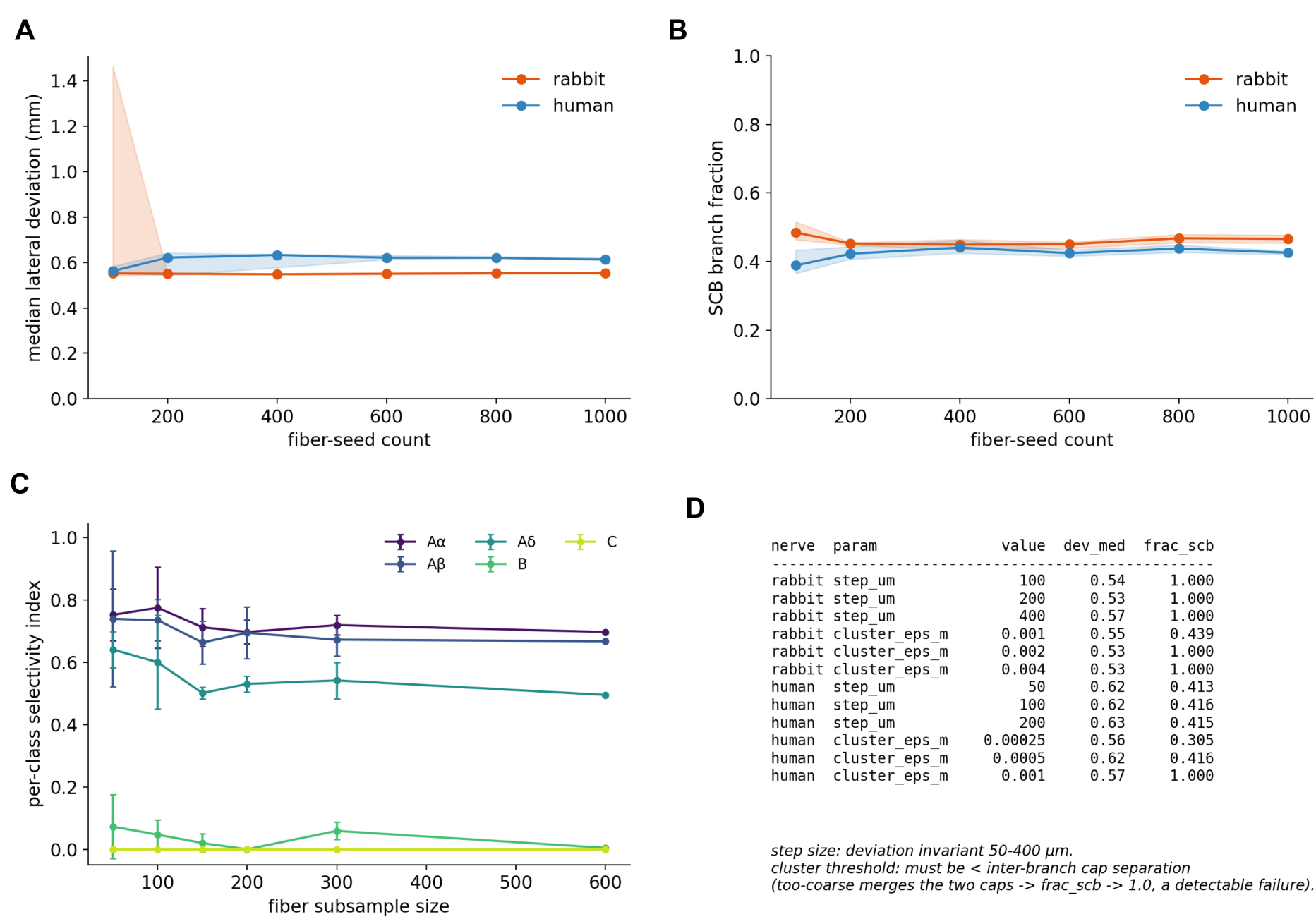
Convergence and robustness of the streamline trajectory and branch-detection step. **A:** Median fiber lateral deviation from the straight chord versus fiber-seed count (three random seedings per count; band is the realization range), rabbit and human. **B:** Fraction of fibers assigned to the superior cardiac branch versus seed count (line is the mean; band the realization range). **C:** Per-class Veraart selectivity index versus random fiber-subsample size (mean ± range over three realizations; human), showing the selectivity estimate is converged. **D:** Sensitivity of the median deviation and branch fraction to the streamline integration step size and the cap-clustering threshold: the step is immaterial over 50–400 µm, whereas the clustering threshold must be set below the inter-branch cap separation (a too-coarse threshold merges the two caps—a detectable failure mode).

**S12 Fig.**
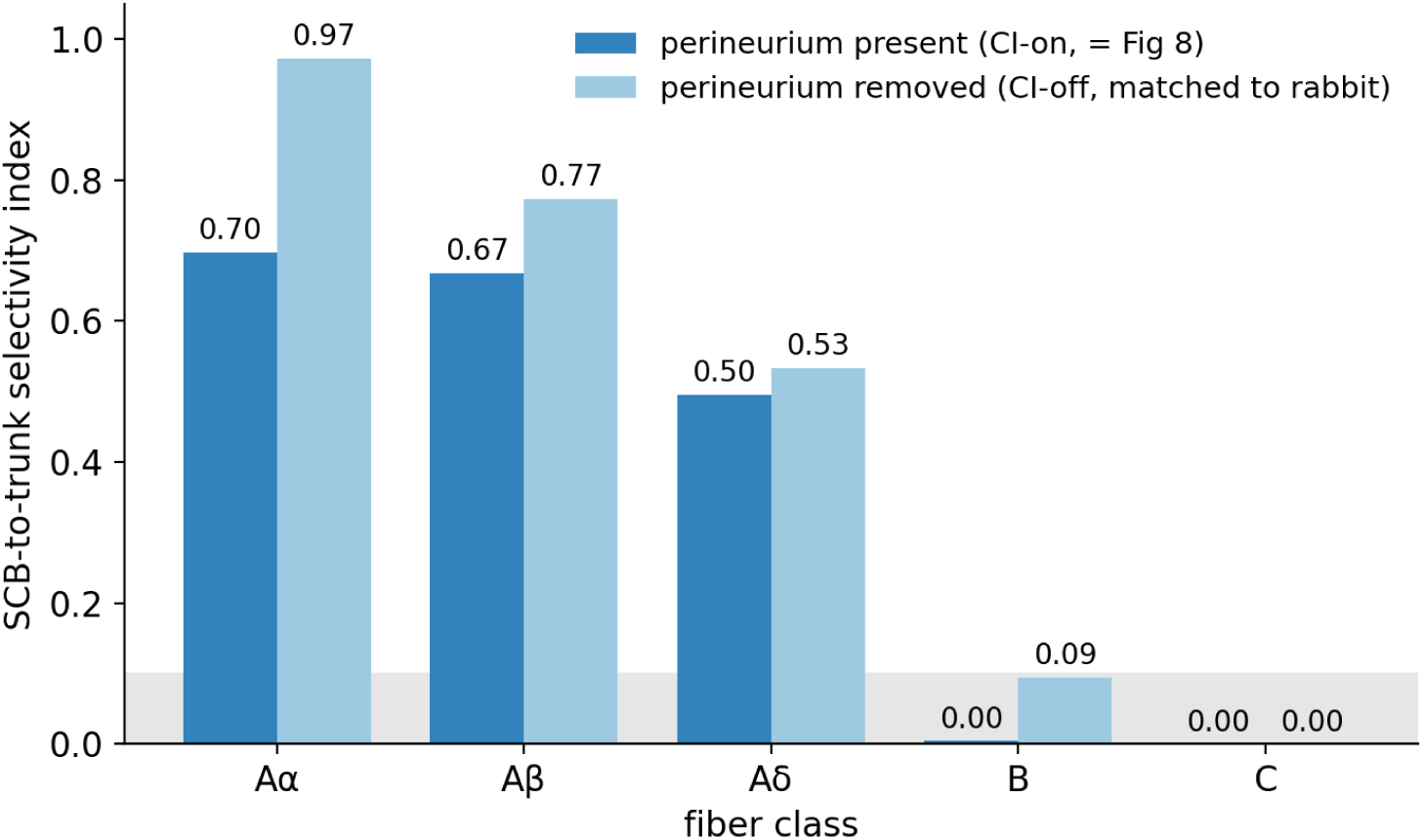
Matched-perineurium control (human cervical vagus). Per-class SCB-to-trunk Veraart selectivity index with the perineurium contact-impedance sheet present (CI-on, identical to Fig 8) versus removed (CI-off, matching the rabbit’s no-perineurium treatment), on the same mesh, fixed longitudinal tripole, and 600-fiber population (only the boundary condition differs). Removing the perineurium sharpens large-fiber selectivity but leaves the small cardiac B and unmyelinated C fibers non-separable (shaded band): the perineurium treatment shifts any per-class index by at most 0.28 (0.09 for B), far below the 0.74 rabbit–human B-fiber gap, so the rabbit-versus-human contrast is anatomical rather than an artifact of perineurium treatment.

**S13 Fig.**
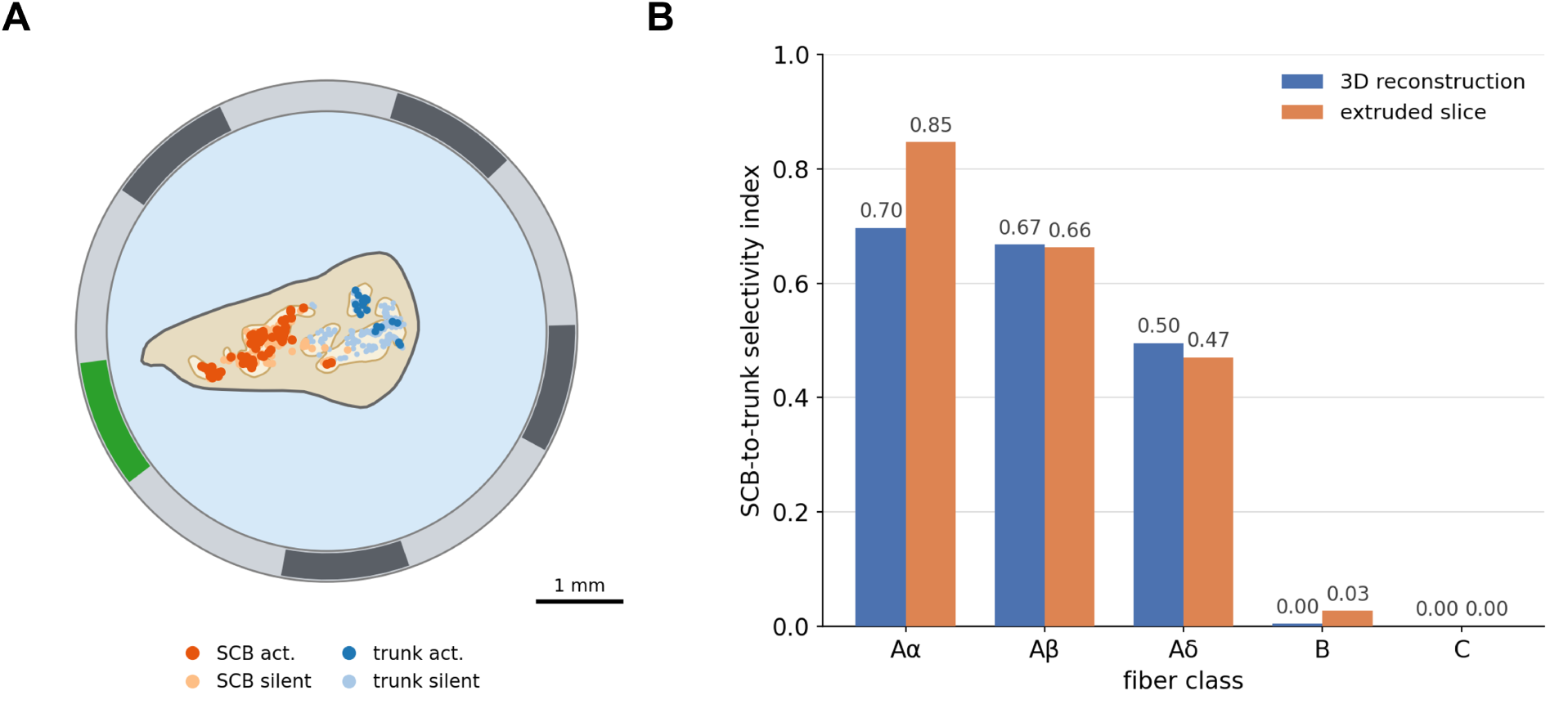
Extruded-slice approximation versus full three-dimensional reconstruction (human cervical vagus). **A:** the extruded counterpart of the reconstructed human nerve—its cuff-plane cross-section (dashed cuff lumen; five fascicle outlines) extruded into straight, constant-section tubes—with straight fibers seeded at the same cuff-plane positions as the three-dimensional fibers and carrying the same branch labels (red, superior cardiac branch; blue, trunk continuation; 281 versus 319 fibers). The longitudinal guarded tripole (cathode 13, guard anodes 8 and 18) occupies a single azimuthal column, marked. **B:** per-class SCB-to-trunk Veraart selectivity index for the same tripole, computed identically for the full three-dimensional reconstruction (Fig 8) and its extruded counterpart under identical 20-contact array, tripole, perineurium contact impedance, fiber population, and 300 µs pulse—so that only the nerve trajectory geometry (curved/branching versus straight extrusion) differs. The approximation agrees within 0.03 for A*β*, A*δ*, B, and C, but overestimates A*α* selectivity by 0.15 (0.85 versus 0.70), the straight extrusion being optimistic about large-fiber separability.

**S14 Fig.**
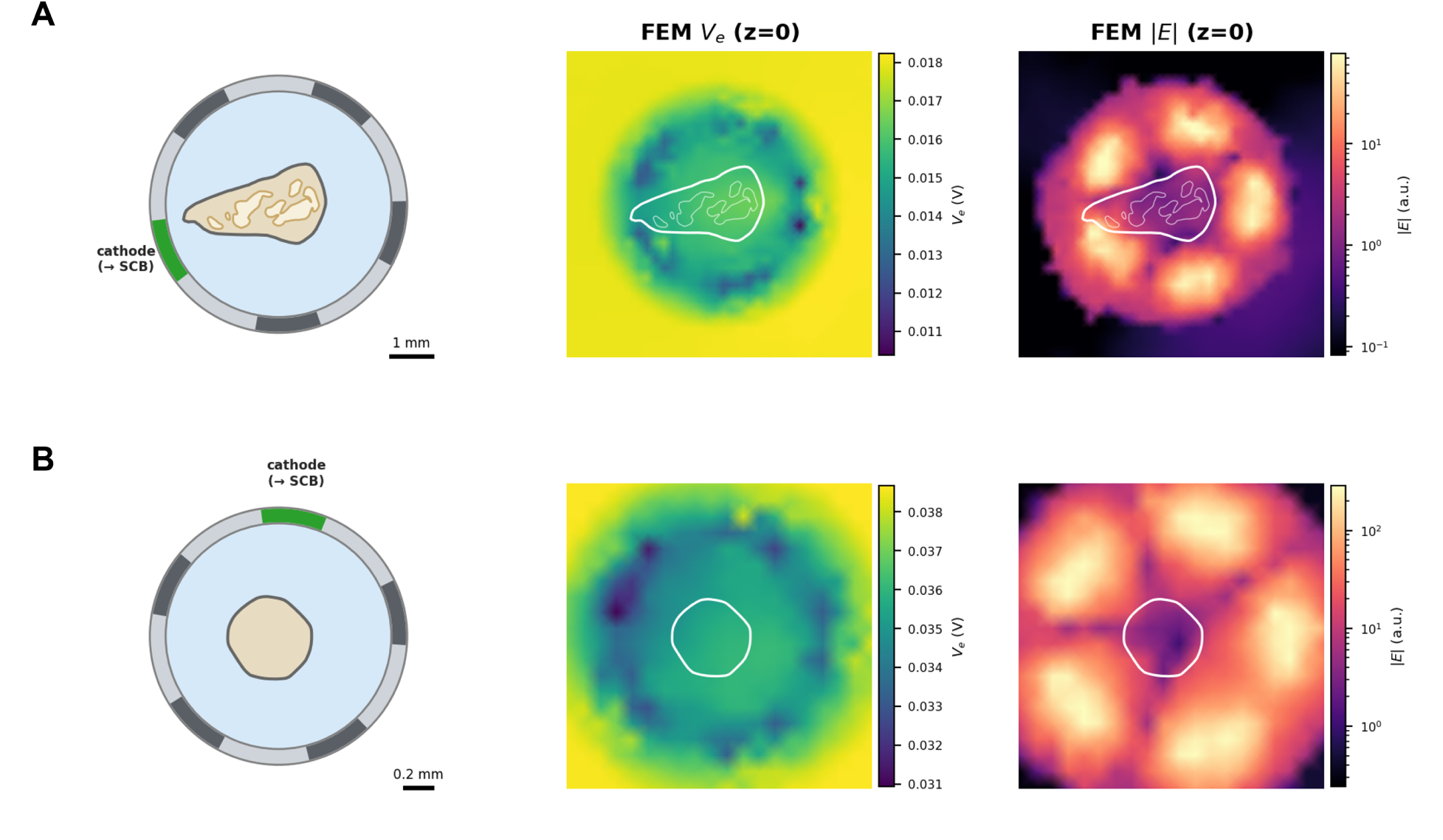
Reconstructed 3D nerves: cuff-plane anatomy and finite-element field. The two micro-CT-reconstructed nerves of this study. Each panel shows, from left to right, the cuff cross-section (saline lumen, silicone cuff, epineurium, and fascicles), the finite-element extracellular potential *V*_e_, and the electric-field magnitude |*E*| on the cuff cross-section (*z* = 0). In the cross-section, the 20-contact array appears as its five azimuthal columns—one axial row in plane—with the cathode column toward the superior cardiac branch in green and the longitudinal-tripole guards in the out-of-plane rows above and below. **(A)** Human nerve (Fig 8): five fascicles, cuff inner radius 1.99 mm. **(B)** Rabbit nerve (Fig 7): monofascicular, cuff inner radius 0.73 mm. Colour scales are per nerve, since the two differ roughly threefold in radius.

**S15 Fig.**
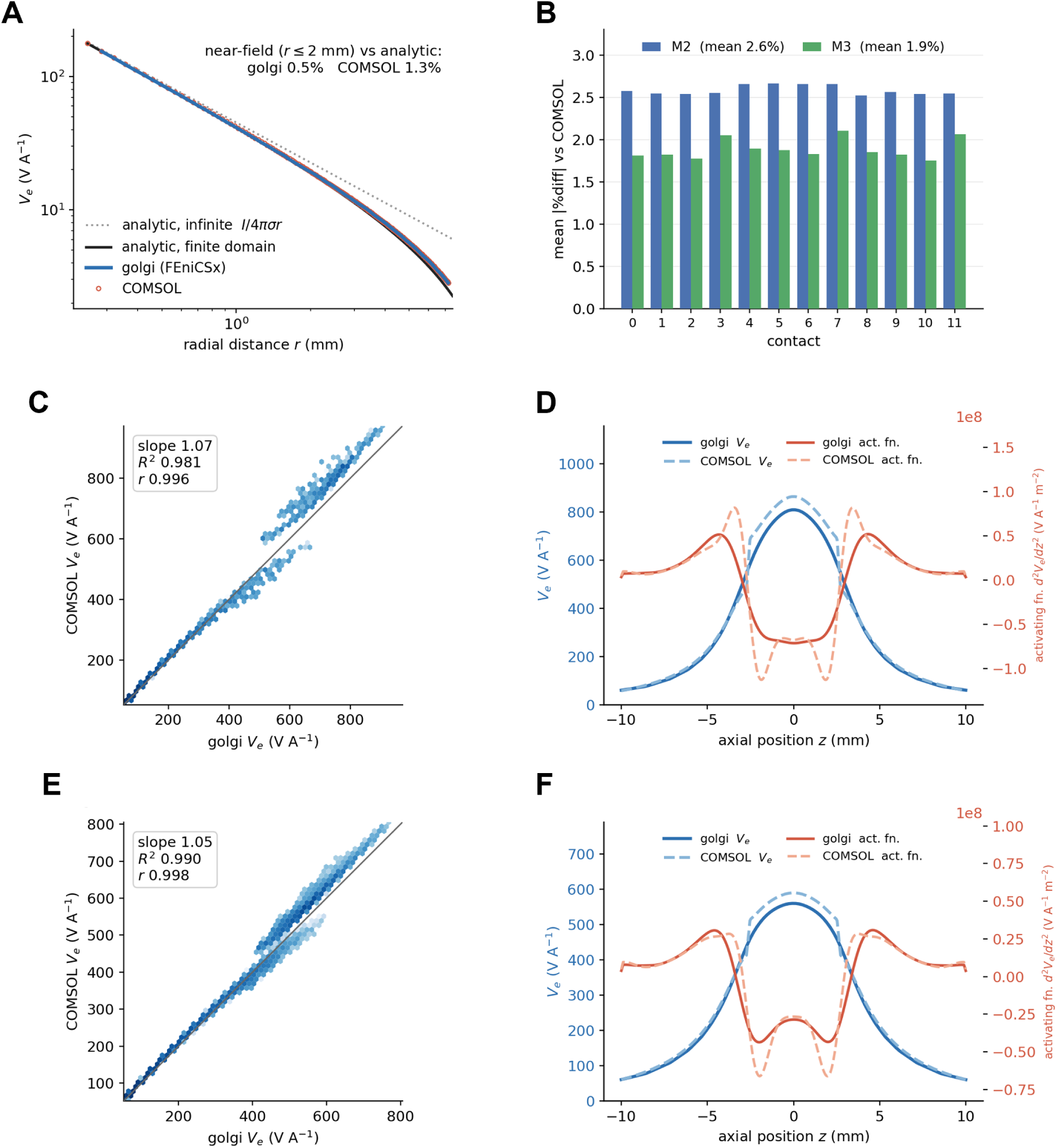
Cross-validation of the open FEniCSx solver against COMSOL. Three COMSOL reference models, built and meshed independently at the finest mesh setting. **A:** M1, a current monopole in a grounded saline cylinder—golgi (FEniCSx) and COMSOL extracellular potential *V_e_*(*r*) versus the finite-domain analytic solution 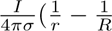 (the infinite-medium *I/*4*πσr* shown dotted for reference); both solvers reproduce the analytic potential to a near/mid-field mean (*r* ≤ 2 mm) of 0.5 % (golgi) and 1.3 % (COMSOL), and golgi is mesh-converged across three mesh levels. **B:** mean absolute per-contact lead-field difference (normalised by each contact’s peak |*V_e_*|) across the 12 contacts of the two multifascicular models. **C–F:** for each multifascicular nerve, the golgi-versus-COMSOL point-for-point density scatter (left; 12 contacts, ∼10^5^ sample positions; slope, *R*^2^, Pearson *r*) and the centre-contact *V_e_*(*z*) profile with its activating function *d*^2^*V_e_/dz*^2^ along the longest fiber (right). **C,D:** M2, an idealized cuff with seven fascicles under full physics (anisotropic endoneurium + perineurium contact impedance), the geometry rebuilt independently in COMSOL. **E,F:** M3, a real swine cervical vagus (sub-4/sam-3, 40 fascicles). golgi reproduces the commercial solver to a mean per-contact difference of 2.6 % (M2) and 1.9 % (M3), *R*^2^ ≥ 0.98, confirming that the open FEniCSx stack—including the anisotropic conductivity and the perineurium contact-impedance boundary—reproduces the COMSOL reference.

**S1 Table.** Nerve sample cohort. Per-sample geometry (fascicle count, dimensions) and fiber-count summary for all 36 modeled nerves (18 swine, 18 human), drawn from the SPARC *Quantified vagus nerve morphology across species* resource—the pig [51] and human [52] datasets of Pelot et al. [7]. *r*_max_ is the maximum endoneurial radius within the cuff; “On-mesh” is the fraction of fiber sample points contained in the solved finite-element domain. Sample IDs abbreviate the SPARC subject/sample designation (e.g., 10/1 = sub-10, sam-1).

| Sample | Species | Fascicles | $r_{\max}$ (mm) | Tets | Fibers | Perineurium ($\mu\text{m}$) | On-mesh |
| --- | --- | --- | --- | --- | --- | --- | --- |
| 46/2 | human | 2 | 1.19 | 1,523,554 | 1000 | 87.2 | 100% |
| 47/2 | human | 5 | 1.03 | 1,450,880 | 1000 | 75.0 | 98% |
| 50/2 | human | 15 | 1.44 | 2,005,779 | 1000 | 51.6 | 100% |
| 53/2 | human | 5 | 1.11 | 1,506,513 | 1001 | 300.3 | 100% |
| 54/2 | human | 7 | 0.86 | 1,317,692 | 1000 | 33.2 | 100% |
| 54/3 | human | 8 | 0.96 | 1,446,581 | 999 | 122.2 | 100% |
| 55/3 | human | 1 | 0.55 | 980,754 | 1000 | 48.0 | 100% |
| 56/1 | human | 6 | 0.89 | 1,320,094 | 999 | 19.6 | 100% |
| 56/3 | human | 5 | 1.01 | 1,435,248 | 1000 | 56.2 | 100% |
| 57/1 | human | 3 | 0.78 | 1,194,232 | 1000 | 22.2 | 100% |
| 57/3 | human | 7 | 1.03 | 1,469,387 | 1000 | 25.2 | 100% |
| 58/1 | human | 13 | 0.80 | 1,370,692 | 1000 | 15.1 | 100% |
| 58/3 | human | 12 | 1.41 | 1,907,222 | 1000 | 49.1 | 100% |
| 63/1 | human | 2 | 0.74 | 1,142,403 | 1000 | 40.4 | 100% |
| 64/1 | human | 1 | 1.13 | 1,440,275 | 1000 | 35.0 | 100% |
| 65/1 | human | 2 | 0.91 | 1,280,942 | 1000 | 39.3 | 100% |
| 69/1 | human | 8 | 1.28 | 1,701,459 | 1000 | 33.9 | 100% |
| 70/1 | human | 2 | 1.10 | 1,440,155 | 1000 | 24.7 | 100% |
| 10/1 | swine | 27 | 1.00 | 1,779,435 | 997 | 7.4 | 99% |
| 11/1 | swine | 39 | 1.14 | 2,165,952 | 999 | 8.9 | 99% |
| 11/3 | swine | 34 | 0.85 | 1,792,386 | 999 | 7.2 | 100% |
| 12/1 | swine | 52 | 1.21 | 2,418,124 | 1001 | 8.6 | 99% |
| 12/3 | swine | 34 | 0.80 | 1,762,503 | 1001 | 6.8 | 99% |
| 13/1 | swine | 63 | 1.18 | 2,640,922 | 999 | 8.0 | 100% |
| 13/3 | swine | 39 | 0.86 | 1,946,475 | 999 | 7.0 | 97% |
| 14/2 | swine | 35 | 1.21 | 2,143,932 | 1001 | 8.1 | 98% |
| 14/3 | swine | 46 | 0.93 | 2,087,844 | 1002 | 6.5 | 100% |
| 15/2 | swine | 37 | 1.22 | 2,162,928 | 994 | 8.8 | 100% |
| 15/3 | swine | 32 | 0.90 | 1,802,965 | 1001 | 7.4 | 100% |
| 4/3 | swine | 38 | 1.18 | 2,173,868 | 998 | 7.4 | 100% |
| 5/1 | swine | 31 | 1.13 | 1,993,476 | 1001 | 8.5 | 99% |
| 5/3 | swine | 32 | 1.00 | 1,908,696 | 1003 | 6.7 | 100% |
| 6/7 | swine | 43 | 1.50 | 2,496,847 | 999 | 9.1 | 100% |
| 8/1 | swine | 27 | 0.98 | 1,787,549 | 998 | 7.3 | 100% |
| 8/7 | swine | 45 | 1.26 | 2,326,999 | 1001 | 8.3 | 100% |
| 9/3 | swine | 57 | 1.32 | 2,596,653 | 1003 | 8.1 | 100% |
| **Total** | 18 swine, 18 human |  |  |  | **35,995** |  |  |

## References

1. González HFJ, Yengo-Kahn A, Englot DJ. Vagus-nerve stimulation for the treatment of epilepsy. Neurosurgery clinics of North America. 2019;1(8):477–82. doi:10.1016/j.nec.2018.12.005.

2. Conway CR, Aaronson S, Sackeim H, George M, Zajecka J, Bunker M, et al. Vagus nerve stimulation in treatment-resistant depression: a one-year, randomized, sham-controlled trial. Brain Stimulation. 2024;18(3):676–89. doi:10.1016/j.brs.2024.12.1191.

3. Tesser JRP, Crowley AR, Box EJ, June JP, Wickersham PB, Valenzuela GJ, et al. Vagus nerve-mediated neuroimmune modulation for rheumatoid arthritis: a pivotal randomized controlled trial. Nature Medicine. 2025;32(1):369–78. doi:10.1038/s41591-025-04114-7.

4. Paton JFR, Z era T, Vadigepalli R, Herring N, Paterson DJ. Multimodal, device-based therapeutic targeting of the cardiovascular autonomic nervous system. Nature Reviews Cardiology. 2025;23(4):255–78. doi:10.1038/s41569-025-01212-4.

5. Navarro X, Krüger TB, Lago N, Micera S, Stieglitz T, Dario P. A critical review of interfaces with the peripheral nervous system for the control of neuroprostheses and hybrid bionic systems. Journal of the Peripheral Nervous System. 2005;10(3):229–58. doi:10.1111/j.1085-9489.2005.10303.x.

6. Grill WM, Norman SE, Bellamkonda RV. Implanted neural interfaces: biochallenges and engineered solutions. Annual Review of Biomedical Engineering. 2009;11(1):1–24. doi:10.1146/annurev-bioeng-061008-124927.

7. Pelot NA, Goldhagen GB, Cariello JE, Musselman ED, Clissold KA, Ezzell JA, et al. Quantified morphology of the cervical and subdiaphragmatic vagus nerves of human, pig, and rat. Frontiers in Neuroscience. 2020;14:1148. doi:10.3389/fnins.2020.601479.

8. Musselman ED, Cariello JE, Grill WM, Pelot NA. ASCENT (Automated Simulations to Characterize Electrical Nerve Thresholds): a pipeline for sample-specific computational modeling of electrical stimulation of peripheral nerves. PLoS Comput Biol. 2021;17(9):e1009285. doi:10.1371/journal.pcbi.1009285.

9. Couppey T, Regnacq L, Giraud R, Romain O, Bornat Y, Kolbl F. NRV: an open framework for in silico evaluation of peripheral nerve electrical stimulation strategies. PLoS Comput Biol. 2024;20(7):e1011826. doi:10.1371/journal.pcbi.1011826.

10. McNeal DR. Analysis of a model for excitation of myelinated nerve. IEEE Transactions on Biomedical Engineering. 1976;BME-23(4):329–37. doi:10.1109/TBME.1976.324593.

11. McIntyre CC, Richardson AG, Grill WM. Modeling the excitability of mammalian nerve fibers: influence of afterpotentials on the recovery cycle. Journal of Neurophysiology. 2002;87(2):995–1006. doi:10.1152/jn.00353.2001.

12. Rattay F. Analysis of models for extracellular fiber stimulation. IEEE Transactions on Biomedical Engineering. 1989;36(7):676–82. doi:10.1109/10.32099.

13. Sweeney JD, Mortimer JT, Durand DM. Modeling of Mammalian Myelinated Nerve for Functional Neuromuscular Stimulation. In: Proceedings of the 9th Annual Conference of the IEEE Engineering in Medicine and Biology Society. Boston, Massachusetts, USA; 1987. p. 1577–8.

14. Sundt D, Gamper N, Jaffe DB. Spike propagation through the dorsal root ganglia in an unmyelinated sensory neuron: a modeling study. Journal of Neurophysiology. 2015;114(6):3140–53. doi:10.1152/jn.00226.2015.

15. Tigerholm J, Petersson M, Obreja O, Lampert A, Carr R, Schmelz M, et al. Modeling Activity-Dependent Changes of Axonal Spike Conduction in Primary Afferent C-Nociceptors. Journal of Neurophysiology. 2014;111(9):1721–35. doi:10.1152/jn.00777.2012.

16. Rattay F, Aberham M. Modeling Axon Membranes for Functional Electrical Stimulation. IEEE Transactions on Biomedical Engineering. 1993;40(12):1201–9. doi:10.1109/10.250575.

17. Schild JH, Clark JW, Hay M, Mendelowitz D, Andresen MC, Kunze DL. A- and C-Type Rat Nodose Sensory Neurons: Model Interpretations of Dynamic Discharge Characteristics. Journal of Neurophysiology. 1994;71(6):2338–58. doi:10.1152/jn.1994.71.6.2338.

18. Schild JH, Kunze DL. Experimental and Modeling Study of Na^+^ Current Heterogeneity in Rat Nodose Neurons and Its Impact on Neuronal Discharge. Journal of Neurophysiology. 1998;78(6):3198–209. doi:10.1152/jn.1997.78.6.3198.

19. Thio BJ, Titus ND, Pelot NA, Grill WM. Reverse-engineered models reveal differential membrane properties of autonomic and cutaneous unmyelinated fibers. PLoS Comput Biol. 2024;20(10):e1012475. doi:10.1371/journal.pcbi.1012475.

20. Lubba C, Guen YL, Jarvis S, Jones NS, Cork S, Eftekhar A, et al. PyPNS: multiscale simulation of a peripheral nerve in Python. Neuroinformatics. 2018;17(1):63–81. doi:10.1007/s12021-018-9383-z.

21. Eiber CD, Payne SC, Biscola NP, Havton LA, Keast JR, Osborne PB, et al. Computational modelling of nerve stimulation and recording with peripheral visceral neural interfaces. Journal of Neural Engineering. 2021;18(6):066020. doi:10.1088/1741-2552/ac36e2.

22. Marshall DP, Farah ES, Musselman ED, Pelot NA, Grill WM. PyFibers: An open-source NEURON-Python package to simulate responses of model nerve fibers to electrical stimulation. PLoS Comput Biol. 2025 12;21(12):e1013764. Available from: 10.1371/journal.pcbi.1013764. doi:10.1371/journal.pcbi.1013764.

23. Hussain MA, Grill WM, Pelot NA. Highly efficient modeling and optimization of neural fiber responses to electrical stimulation. Nature Communications. 2024;15(1):7597. doi:10.1038/s41467-024-51709-8.

24. Ravi N, Gabeur V, Hu YT, Hu R, Ryali CK, Ma T, et al. SAM 2: segment anything in images and videos. International Conference on Learning Representations. 2024. doi:10.48550/arXiv.2408.00714.

25. Ma J, Wang B. Segment anything in medical images. arXivorg. 2023;15:654. doi:10.48550/arXiv.2304.12306.

26. Si H. TetGen, a Delaunay-based quality tetrahedral mesh generator. ACM Transactions on Mathematical Software. 2015;41(2):1–36. doi:10.1145/2629697.

27. Geuzaine C, Remacle JF. Gmsh: a 3-D finite element mesh generator with built-in pre- and post-processing facilities. International Journal for Numerical Methods in Engineering. 2009;79(11):1309–31. doi:10.1002/nme.2579.

28. Bossetti CA, Birdno MJ, Grill WM. Analysis of the quasi-static approximation for calculating potentials generated by neural stimulation. Journal of Neural Engineering. 2008;5(1):44–53. doi:10.1088/1741-2560/5/1/005.

29. Baratta IA, Dean JP, Dokken JS, Habera M, Hale JS, Richardson CN, et al. DOLFINx: the next generation FEniCS problem solving environment. Zenodo. 2023. doi:10.5281/zenodo.10447666.

30. Pelot N, Behrend CE, Grill W. On the parameters used in finite element modeling of compound peripheral nerves. Journal of Neural Engineering. 2018;16(1):016007. doi:10.1088/1741-2552/aaeb0c.

31. Ranck Jr JB, BeMent SL. The specific impedance of the dorsal columns of cat: an anisotropic medium. Experimental Neurology. 1965;11(4):451–63. doi:10.1016/0014-4886(65)90059-2.

32. Weerasuriya A, Spangler RA, Rapoport SI, Taylor RE. AC impedance of the perineurium of the frog sciatic nerve. Biophysical Journal. 1984;46(2):167–74. doi:10.1016/S0006-3495(84)84009-6.

33. Grill WM, Mortimer JT. Electrical properties of implant encapsulation tissue. Annals of Biomedical Engineering. 2006;22(1):23–33. doi:10.1007/BF02368219.

34. Gielen F, Jonge W, Albers BA, Boon K. Electrical conductivity of skeletal muscle tissue: experimental results from different muscles in vivo. Medical amp; Biological Engineering amp; Computing. 1984;22(6):569–77. doi:10.1016/0303-8467(84)90218-X.

35. Horch KW, Dhillon G. Neuroprosthetics: Theory and Practice. 2nd ed. Horch KW, Kipke DR, editors. Singapore: World Scientific; 2004. doi:10.1142/10368.

36. Kluge C. Fundamentals of Materials Science and Engineering: An Integrated Approach. 4th ed. Hoboken, NJ: John Wiley & Sons; 2016.

37. Podesta M. Understanding the Properties of Matter. 2nd ed. London: Taylor & Francis; 1996. doi:10.4324/9780203450611.

38. Marshall DP, Upadhye AR, Buyukcelik ON, Shoffstall AJ, Grill WM, Pelot NA. Computational modeling of human vagus nerve stimulation with three-dimensional fascicular morphology. APL Bioengineering. 2026;10(1):016112. doi:10.1063/5.0308450.

39. Marshall DP, Farah ES, Musselman ED, Pelot NA, Grill WM. PyFibers; 2026. Software. Zenodo. Available from: 10.5281/zenodo.19595192. doi:10.5281/zenodo.19595192.

40. Hines ML, Carnevale NT. The NEURON simulation environment. Neural Computation. 1997;9(6):1179–209. doi:10.1162/neco.1997.9.6.1179.

41. Hussain MA. wmglab-duke/axonml: v1.0.2; 2024. Software. Zenodo. Available from: 10.5281/zenodo.12752386. doi:10.5281/zenodo.12752386.

42. Hursh JB. Conduction velocity and diameter of nerve fibers. American Journal of Physiology-Legacy Content. 1939;127(1):131–9. doi:10.1152/ajplegacy.1939.127.1.131.

43. Weiss G. Sur la possibilité de rendre comparables entre eux les appareils servant ‘a l’excitation électrique. Archives Italiennes de Biologie. 1990;35:413–46. doi:10.4449/AIB.V35I1.1355.

44. Lapicque L. Recherches quantitatives sur l’excitation électrique des nerfs traitée comme une polarisation. Journal de Physiologie et de Pathologie Géńerale. 1907;9:620–35.

45. Yoo PB, Lubock NB, Hincapie JG, Ruble SB, Hamann JJ, Grill WM. High-resolution measurement of electrically-evoked vagus nerve activity in the anesthetized dog. Journal of Neural Engineering. 2013;10(2):026003. doi:10.1088/1741-2560/10/2/026003.

46. Nannini N, Horch K. Muscle recruitment with intrafascicular electrodes. IEEE Transactions on Biomedical Engineering. 1991;38(8):769–76. doi:10.1109/10.83589.

47. Yoshida K, Horch K. Selective stimulation of peripheral nerve fibers using dual intrafascicular electrodes. IEEE Transactions on Biomedical Engineering. 1993;40(5):492–4. doi:10.1109/10.243412.

48. Bucksot JE, Wells AJ, Rahebi KC, Sivaji V, Romero-Ortega M, Kilgard MP, et al. Flat electrode contacts for vagus nerve stimulation. PLOS ONE. 2019;14(11):e0215191. doi:10.1371/journal.pone.0215191.

49. Kronsteiner B, Carrero-Rojas G, Reissig LF, Moghaddam AS, Schwendt KM, Gerges S, et al. Characterization, number, and spatial organization of nerve fibers in the human cervical vagus nerve and its superior cardiac branch. Brain Stimulation. 2024;17(3):510–24. doi:10.1016/j.brs.2024.04.016.

50. Veraart C, Grill WM, Mortimer JT. Selective control of muscle activation with a multipolar nerve cuff electrode. IEEE Transactions on Biomedical Engineering. 1993;40(7):640–53. doi:10.1109/10.237694.

51. Pelot NA, Goldhagen GB, Cariello JE, Grill WM. Quantified morphology of the pig vagus nerve. SPARC Portal; 2020. Dataset, Quantified vagus nerve morphology across species (Version 4), RRID:SCR 017041. doi:10.26275/maq2-eii4.

52. Pelot NA, Ezzell JA, Goldhagen GB, Cariello JE, Clissold KA, Grill WM. Quantified morphology of the human vagus nerve with anti-claudin-1. SPARC Portal; 2025. Dataset, Quantified vagus nerve morphology across species (Version 8), RRID:SCR 017041. doi:10.26275/qgja-q7zd.

53. Haberbusch M, Lung D, Jia Y, Blumer R, Reissig LF, Zopf LM, et al.. Rabbit Cervical Vagus Nerve Micro-Computed Tomography Dataset from Nodose Ganglion to Superior Cardiac Branch with Tissue Segmentations and 3D Reconstruction; 2026. Dataset. Zenodo. Available from: 10.5281/zenodo.21006413. doi:10.5281/zenodo.21006413.

54. Haberbusch M, Zopf LM, Heimel P, Blumer R, Reissig LF. Human Cervical Vagus Nerve µCT Dataset from Nodose Ganglion to Superior Cardiac Branch with Tissue Segmentations and 3D Reconstruction; 2026. Dataset. Zenodo. Available from: 10.5281/zenodo.21007683. doi:10.5281/zenodo.21007683.

55. Haberbusch M. Reproduction bundles for the golgi peripheral nerve stimulation modeling platform; 2026. Dataset. Zenodo. Available from: 10.5281/zenodo.21300945. doi:10.5281/zenodo.21300945.

56. Choi AQ, Cavanaugh JK, Durand DM. Selectivity of multiple-contact nerve cuff electrodes: a simulation analysis. IEEE Transactions on Biomedical Engineering. 2001;48(2):165–72. doi:10.1109/10.909637.

57. Ardell JL, Nier H, Hammer M, Southerland EM, Ardell CL, Beaumont E, et al. Defining the neural fulcrum for chronic vagus nerve stimulation: implications for integrated cardiac control. Journal of Physiology. 2017;595(22):6887–903. doi:10.1113/JP274678.

58. Wilkinson MD, Dumontier M, Aalbersberg IJ, Appleton G, Axton M, Baak A, et al. The FAIR Guiding Principles for scientific data management and stewardship. Scientific Data. 2016;3(1):160018. doi:10.1038/sdata.2016.18.

59. Haberbusch M, Lung D, Jia Y, Fachino M, Moro A. golgi: an open platform for image-to-recruitment modeling of peripheral nerve stimulation; 2026. Software. Zenodo. Available from: 10.5281/zenodo.21301866. doi:10.5281/zenodo.21301866.

